# Single cell analysis reveals the impact of age and maturation stage on the human oocyte transcriptome

**DOI:** 10.1101/2020.09.25.309658

**Authors:** Silvia Llonch, Montserrat Barragán, Paula Nieto, Anna Mallol, Marc Elosua-Bayes, Patricia Lorden, Sara Ruiz, Filippo Zambelli, Holger Heyn, Rita Vassena, Bernhard Payer

**Affiliations:** Centre for Genomic Regulation (CRG), The Barcelona Institute of Science and Technology, Dr. Aiguader 88, 08003 Barcelona, Spain; Clínica EUGIN, Barcelona, Spain; CNAG-CRG, Barcelona, Spain; Universitat Pompeu Fabra (UPF), Barcelona, Spain

**Keywords:** human oocyte, ageing, fertility, advanced maternal age, assisted reproduction, transcriptomics, single cell RNA-Seq

## Abstract

**Study question:** To which degree does maternal age affect the transcriptome of human oocytes at the germinal vesicle (GV) stage or at metaphase II after maturation *in vitro* (IVM-MII)?

**Summary answer:** While the oocytes’ transcriptome is predominantly determined by maturation stage, transcript levels of genes related to chromosome segregation, mitochondria and RNA processing are affected by age after *in vitro* maturation of denuded oocytes.

**What is known already:** Female fertility is inversely correlated with maternal age due to both a depletion of the oocyte pool and a reduction in oocyte developmental competence. Few studies have addressed the effect of maternal age on the human mature oocyte (MII) transcriptome, which is established during oocyte growth and maturation, and the pathways involved remain unclear. Here, we characterize and compare the transcriptomes of a large cohort of fully grown GV and IVM-MII oocytes from women of varying reproductive age.

**Study design, size, duration:** In this prospective molecular study, 37 women were recruited from May 2018 to June 2019. The mean age was 28.8 years (SD=7.7, range 18-43). A total of 72 oocytes were included in the study at GV stage after ovarian stimulation, and analyzed as GV (n=40) and *in vitro* matured oocytes (IVM-MII; n=32).

**Participants/materials, setting, methods:** Denuded oocytes were included either as GV at the time of ovum pick-up or as IVM-MII after *in vitro* maturation for 30 hours in G2^™^ medium, and processed for transcriptomic analysis by single-cell RNA-seq using the Smart-seq2 technology. Cluster and maturation stage marker analysis were performed using the Seurat R package. Genes with an average fold change greater than 2 and a p-value < 0.01 were considered maturation stage markers. A Pearson correlation test was used to identify genes whose expression levels changed progressively with age. Those genes presenting a correlation value (R) >= |0.3| and a p-value < 0.05 were considered significant.

**Main results and the role of chance:** First, by exploration of the RNA-seq data using tSNE dimensionality reduction, we identified two clusters of cells reflecting the oocyte maturation stage (GV and IVM-MII) with 4,445 and 324 putative marker genes, respectively. Next we identified genes, for which RNA levels either progressively increased or decreased with age. This analysis was performed independently for GV and IVM-MII oocytes. Our results indicate that the transcriptome is more affected by age in IVM-MII oocytes (1,219 genes) than in GV oocytes (596 genes). In particular, we found that genes involved in chromosome segregation and RNA splicing significantly increase in transcript levels with age, while genes related to mitochondrial activity present lower transcript levels with age. Gene regulatory network analysis revealed potential upstream master regulator functions for genes whose transcript levels present positive (*GPBP1, RLF, SON, TTF1*) or negative (*BNC1, THRB*) correlation with age.

**Limitations, reasons for caution:** IVM-MII oocytes used in this study were obtained after *in vitro* maturation of denuded GV oocytes, therefore, their transcriptome might not be fully representative of *in vivo* matured MII oocytes.

The Smart-seq2 methodology used in this study detects polyadenylated transcripts only and we could therefore not assess non-polyadenylated transcripts.

**Wider implications of the findings:** Our analysis suggests that advanced maternal age does not globally affect the oocyte transcriptome at GV or IVM-MII stages. Nonetheless, hundreds of genes displayed altered transcript levels with age, particularly in IVM-MII oocytes. Especially affected by age were genes related to chromosome segregation and mitochondrial function, pathways known to be involved in oocyte ageing. Our study thereby suggests that misregulation of chromosome segregation and mitochondrial pathways also at the RNA-level might contribute to the age-related quality decline in human oocytes.

**Study funding/competing interest(s):** This study was funded by the AXA research fund, the European commission, intramural funding of Clinica EUGIN, the Spanish Ministry of Science, Innovation and Universities, the Catalan Agència de Gestió d’Ajuts Universitaris i de Recerca (AGAUR) and by contributions of the Spanish Ministry of Economy, Industry and Competitiveness (MEIC) to the EMBL partnership and to the “Centro de Excelencia Severo Ochoa”.

The authors have no conflict of interest to declare.

## Introduction

During the last decades maternal age at the birth of the first child has significantly increased (Matthews and Hamilton, 2009). Concurrently, assisted reproductive techniques are used with increasing frequency in women older than 35, as natural fertility decreases significantly past this age (Howles *et al*., 2006). Two reasons for this decline in fertility, which are usually concomitant but not necessarily connected, are the depletion of the oocyte pool over time and a reduction in oocyte quality, which leads to increased incidence of aneuploidies, lower embryo developmental rates and increased pregnancy loss (Nagaoka *et al*., 2012).

During fetal development, oocytes initiate meiosis and arrest at the diplotene stage of prophase I (also known as dictyate stage), where the oocytes present a characteristic nucleus called germinal vesicle (GV) and remain quiescent for several years. This meiotic arrest persists until oocytes mature to the metaphase II (MII) stage at which they are ovulated throughout the reproductive lifespan of a woman. Many biochemical pathways have been postulated to explain the oocyte quality drop and the increased incidence of embryo aneuploidies related to ageing (Nagaoka *et al*., 2012). During meiotic arrest, the linkage between chromatids is maintained by crossing overs and proteins such as cohesins. In the mouse model it has been suggested that REC8-Cohesin participates in maintaining chromosome cohesion without turnover in oocytes arrested for months (Burkhardt *et al*., 2016). As well the centromeric histone variant CENP-A, which is important for kinetochore identity and proper chromosome segregation, is incorporated early during mouse oocyte development and remains at centromeres without significant turnover until oocyte maturation much later in life (Smoak *et al*., 2016), although in other species like starfish, CENP-A might be gradually replenished in meiotically arrested oocytes (Swartz *et al*., 2019). It is unclear to which degree proteins like REC8 and CENP-A are turned over in human oocytes over time. Furthermore, studies in mouse and human suggest that age-related chromosome segregation errors could be due to gradual loss of cohesion and kinetochore compaction (Burkhardt *et al*., 2016; Smoak *et al*., 2016; Toth and Jessberger, 2016; Gruhn *et al*., 2019; Zielinska *et al*., 2019) and altered microtubule dynamics leading to non-disjunction events (Nakagawa and FitzHarris, 2017). In addition, mitochondria-related defects (May-Panloup *et al*., 2016; Almansa-Ordonez *et al*., 2020) and epigenetic changes affecting gene expression (Ge *et al*., 2015; Chamani and Keefe, 2019) contribute to the age-related oocyte quality decline.

A deeper understanding of the mechanisms driving the decay of oocyte quality with age is important. While some age-related defects are based on low protein turnover and the associated changes in protein levels over the years during which oocytes remain dormant in meiotic arrest, it is less clear to which degree the transcriptome plays a role in oocyte ageing. Several studies have been performed in human oocytes using microarray analysis (reviewed in (Labrecque and Sirard, 2014). Grøndahl and colleagues found age-related differences in transcript abundance in *in vivo* matured MII oocytes (Grøndahl *et al*., 2010), and analyzed their transcriptome in the quiescent state (Grøndahl *et al*., 2013). Due to the transcriptionally quiescent state of the fully grown GV oocyte, the RNA poly(A) tail length is controlled, resulting in differential transcript-specific RNA stability and translational state (Jalkanen *et al*., 2014). Recent studies applied RNA-seq using oligo(dT)-based reverse transcription to identify differences between GV and MII oocytes from women of varying reproductive age (Reyes *et al*., 2017); (Zhang *et al*., 2020).

Single-cell RNA sequencing (scRNA-seq) techniques developed over the last decade are among the leading tools for exploring tissue heterogeneity at cellular level (Pijuan-Sala *et al*., 2018; Svensson *et al*., 2018). They facilitate the identification of transcriptional differences between cells that would remain undetectable with conventional bulk RNA sequencing techniques. In this study, we apply scRNA-seq analysis on the poly(A)-RNA transcriptome of single human GV oocytes obtained during ovum pick-up after ovarian stimulation of young and advanced maternal age women. Additionally, applying an experimental *in vitro* maturation protocol towards the MII stage, we identified differences in abundance of RNAs related to specific biological processes (chromosome segregation, cell cycle regulation, mitochondrial function and RNA metabolism) that were correlated with the women’s age. Furthermore we found through network analysis potential master regulators involved in reproductive ageing. Thus our data suggests that RNA-turnover might play an instructive role in oocyte ageing thereby advancing our understanding of the reproductive ageing process.

## Materials and Methods

### Ethical approval

Approval to conduct this study was obtained from the Ethics Committee for Clinical Research (CEIm) of Clinica Eugin before the beginning. All women included in the study gave their written informed consent prior to inclusion.

### Study population

A total of 37 women (n=25 oocyte donors and n=12 patients) were included in the study. All women had a body mass index (BMI) <33, normal karyotype and no systemic or reproductive conditions, such as endometriosis. Oocytes from only one cycle of ovarian stimulation per woman were included. For single-oocyte RNA-seq analysis, 72 oocytes were collected; 40 of them were included in the study as GV, while 32 were included as MII after *in vitro* maturation (IVM-MII). The average women age was 28.8 ± 7.7 (range 18-43), with a mean ovarian reserve (measured by antral follicle count; AFC) of 22.1 ± 10.7 (range 4-46). The characteristics of the ovum pick-up for each participant are shown in Supp. Table 1.

### Ovarian stimulation, oocyte retrieval and *in vitro* maturation

Women were stimulated with highly purified urinary hMG (Menopur®, Ferring, Spain) or follitropin alpha (Gonal®, Merck-Serono, Spain), with daily injections of 150-300 IU (Blazquez *et al*., 2014). A GnRH antagonist (0.25 mg of Cetrorelix acetate, Cetrotide®, Merck Serono, Spain) was administered daily from day 6 of stimulation (for donors) or from when a follicle of 14 mm or estradiol ≥ 400 pg/ml was detected (for patients) (Olivennes *et al*., 1996). In the case of donors, when 3 or more follicles of ≥18 mm of diameter were observed, final oocyte maturation was triggered with 0.2 mg of triptorelin (Decapeptyl®, Ipsen Pharma, Spain). In the case of patients, when 3 follicles of ≥17 mm of diameter were observed, final oocyte maturation was triggered with 250 µg of alpha-choriogonadotropin (Ovitrelle®, MERCK) or 0.3 mg of triptorelin (Decapeptyl®, Ipsen Pharma, Spain). Oocyte retrieval was performed 36h later by ultrasound-guided transvaginal follicular aspiration. Oocytes were denuded 30 minutes after pick-up by exposure to 80 IU/ml hyaluronidase (Hyase-10x, Vitrolife, Sweden) in G-MOPS medium (Vitrolife, Sweden), followed by gentle pipetting. Once denuded, oocytes were scored for polar body presence and immature GV were either processed immediately as GV or further cultured *in vitro* in 50 µl of G2-PLUS (Vitrolife, Sweden) medium in a humidified atmosphere of 6%CO2/94% at 37°C for 30 hours, when they were checked again for polar body presence and processed.

### Single-cell RNA sequencing

Full-length single-cell RNA-seq libraries were prepared using the Smart-seq2 protocol (Picelli *et al*., 2013) with minor modifications. Briefly, oocytes were dezoned with Pronase (Roche Diagnostics, Spain), and individually placed in 2.3 µl of a lysis buffer containing 0.2% Triton-X100 (T8787, Sigma) and 1 U/µl RNAse inhibitor (N8080119, Applied Biosystem), and stored at −80°C until use. Reverse transcription was performed using SuperScript II (ThermoFisher Scientific) in the presence of 1 µM oligo-dT_30_VN (IDT), 1 µM template-switching oligonucleotides (QIAGEN), and 1 M betaine. cDNA was amplified using the KAPA Hifi Hotstart ReadyMix (Kapa Biosystems) and IS PCR primer (IDT), with 20 cycles of amplification. Following purification with Agencourt Ampure XP beads (Beckmann Coulter), product size distribution and quantity were assessed on a Bioanalyzer using a High Sensitivity DNA Kit (Agilent Technologies). A total of 140 pg of the amplified cDNA was fragmented using Nextera XT (Illumina) and amplified with Nextera XT indexes (Illumina). Products of each of the 96-well plate were pooled and purified twice with Agencourt Ampure XP beads (Beckmann Coulter). Final libraries were quantified and checked for fragment size distribution using a Bioanalyzer High Sensitivity DNA Kit (Agilent Technologies). Pooled sequencing of Nextera libraries was carried out using a HiSeq4000 (Illumina) to an average sequencing depth of >1 million reads per cell. Sequencing was carried out as paired-end (PE75) reads with library indexes corresponding to cell barcodes (Unique dual indexing).

### Data analysis

#### scRNA-seq initial processing

Raw sequencing data was obtained from the Smart-seq2 protocol as described elsewhere (Guillaumet-Adkins *et al*., 2017) with minor modifications. Briefly, initial quality check on the FASTA files was carried out with FastQC quality control suite. Samples that reached quality standards were processed to deconvolute reads and assign them to a single cell by demultiplexing according to pool barcodes. PolyT reads were removed. Sequencing reads were mapped with the STAR v2.5.4b RNA aligner (Dobin *et al*., 2013) with default parameters and the reference genome was Gencode release 32 (assembly GRCh38.p13). Gene quantification was carried out using UMI to account for amplification biases (allowing an edit distance up to two nucleotides in UMI comparisons). Only unambiguously mapped reads were considered (Guillaumet-Adkins *et al*., 2017). Mapped reads were assigned to genes when they overlapped with exons. Only oocytes with ≥1,000 UMIs, ≥10,000 reads, and ≤30% mitochondrial transcripts (35 GVs and 31 IVM-MIIs) were included for further analysis (Supplementary Fig. S1). These metrics were considered jointly to ensure that the discarded cells with high mitochondrial expression were not metabolically active, but instead low quality damaged cells. UMI counts were normalized using Seurat’s (Butler *et al*., 2018; Stuart *et al*., 2019) SCTransform pipeline, a modeling framework for the normalization and variance stabilization of molecular count data from scRNA-seq (Hafemeister and Satija, 2019), which finds sharper biological differences and avoids most technical/confounding factors compared to Seurat’s standard pipeline.

#### scRNA-seq clustering and differential expression analysis of maturation stages

To cluster the oocytes we i) performed a principal component analysis (PCA) using the scaled and normalized 3,000 most highly variable genes, ii) used the top 20 principal components (PCs) and the *FindNeighbors* function to create a k-nearest neighbor graph based on the lower dimensional embedding, iii) clustered the cells with the *FindClusters* function using the default parameters, including the resolution set to 0.8. To visualise the dataset we used a non-linear dimensionality reduction technique, t-distributed Stochastic Neighbor Embedding (tSNE), on the top 20 PCs and setting the perplexity hyperparameter to 15. To find the cluster markers, in our case corresponding to the maturation stage, the function *FindAllMarkers* was applied, using the “MAST” test (Finak *et al*., 2015) and setting the parameters min.pct and logfc.threshold to 0 and only.pos to TRUE. This analysis was performed using the Seurat *R* package (Butler *et al*., 2018; Stuart *et al*., 2019).

Differentially expressed genes (DEG) presenting an average fold change over 2 between GV and IVM-MII oocytes and a p-value smaller than 0.01 were considered as significant maturation stage markers. EnhancedVolcano R package (Blighe *et al*., 2019) was used for the visualization of the gene’s significance and log2FC (lnFC values obtained from the function *FindAllMarkers* were transformed to log2FC using the formula log2FC = lnFC/ln(2)), the names of the top 10 markers per each maturation stage are plotted.

#### Analysis of gene expression correlation with age

Analysis of gene expression correlation with age was performed using pearson correlation by means of the *cor*.*test* function. This analysis was carried out independently for GV stage oocytes and IVM-MII stage oocytes. Genes presenting a correlation value (R) >= |0.3| and a p-value < 0.05 were considered significantly correlated with maturation stage. Plots of gene expression levels correlating with age were obtained using the ggplot2 R package (Wickham, 2016).

Venn diagrams used to evaluate how many genes correlating with age were shared between the GV stage oocytes and the IVM-MII stage oocytes were plotted using the VennDiagram *R* package (Chen and Boutros, 2011). Lists of the genes positively (increasing RNA-levels with age) or negatively (decreasing RNA-levels with age) correlating with age in GVs and IVM-MIIs were used as an input. Proportional Venn diagrams to intersect maturation stage markers with genes, for which RNA-levels change with age have been generated with the Eulerr *R* package. (Wilkinson, 2012; Micallef and Rodgers, 2014).

#### Gene ontology analysis

GO term enrichment analysis was performed using the *enrichGO* function from the ClusterProfiler *R* package (Shannon *et al*., 2003; Yu *et al*., 2012). The entire list of maturation stage markers (GV and IVM-MII independently), filtered for a significant p-value < 0.01 and a fold change (FC) > |2|, was used as an input. GO terms with an adjusted p-value (FDR method) below 0.05 were considered significant. For GO term enrichment analysis of the genes correlating with age, the function was run four times: genes that positively correlate with age in GVs, genes that negatively correlate with age in GVs and the same for IVM-MIIs. Again, GO terms with an adjusted p-value lower than 0.05 were considered as significant.

#### Gene regulatory network analysis

Gene regulatory networks were analysed using the Genie3 *R* package (Huynh-Thu *et al*., 2010). The normalized expression matrix of genes changing in expression (positive or negative correlation) with age in IVM-MII stage oocytes was given as an input. Either a list of described human transcription factors (TF) (Lambert *et al*., 2018) or all genes changing RNA amount with age were specified in the parameter “regulators” for the Genie3 analysis. The top 2,500 regulatory links were next plotted using Cytoscape3 (Shannon *et al*., 2003). Genes with the highest number of interactions (over 50 for human TF and over 10 for all genes considered as potential regulators and a betweenness centrality value over 0.5 were considered the main regulators of the network. For visualization, the main nodes (blue) size represent the number of interactions with other nodes. The colour green represents genes belonging to GO terms enriched in the list of genes positively correlating (going up in expression) with age in IVM-MII oocytes. The red colour indicates the genes belonging to GO terms enriched in the list of genes negatively correlating (going down in expression) with age in IVM-MII oocytes. Genes with a number of interactions lower than 50 (TF as regulators) or 10 (all genes considered as potential regulators) have a node size equal to 0.

## Results

The main goal of this study was to investigate changes in transcriptome associated with oocyte ageing. For that purpose, we collected GV oocytes from 37 women within an age range of 18-43 years and either subjected them directly at the GV stage (n=40), or after *in vitro* maturation to IVM-MII stage (n=32), to single oocyte RNA-Sequencing using the Smart-seq2 protocol (Picelli *et al*., 2013) (Fig. 1A,B, Supplementary Table 1, Supplementary Fig. S1, S2).

**Figure 1.**
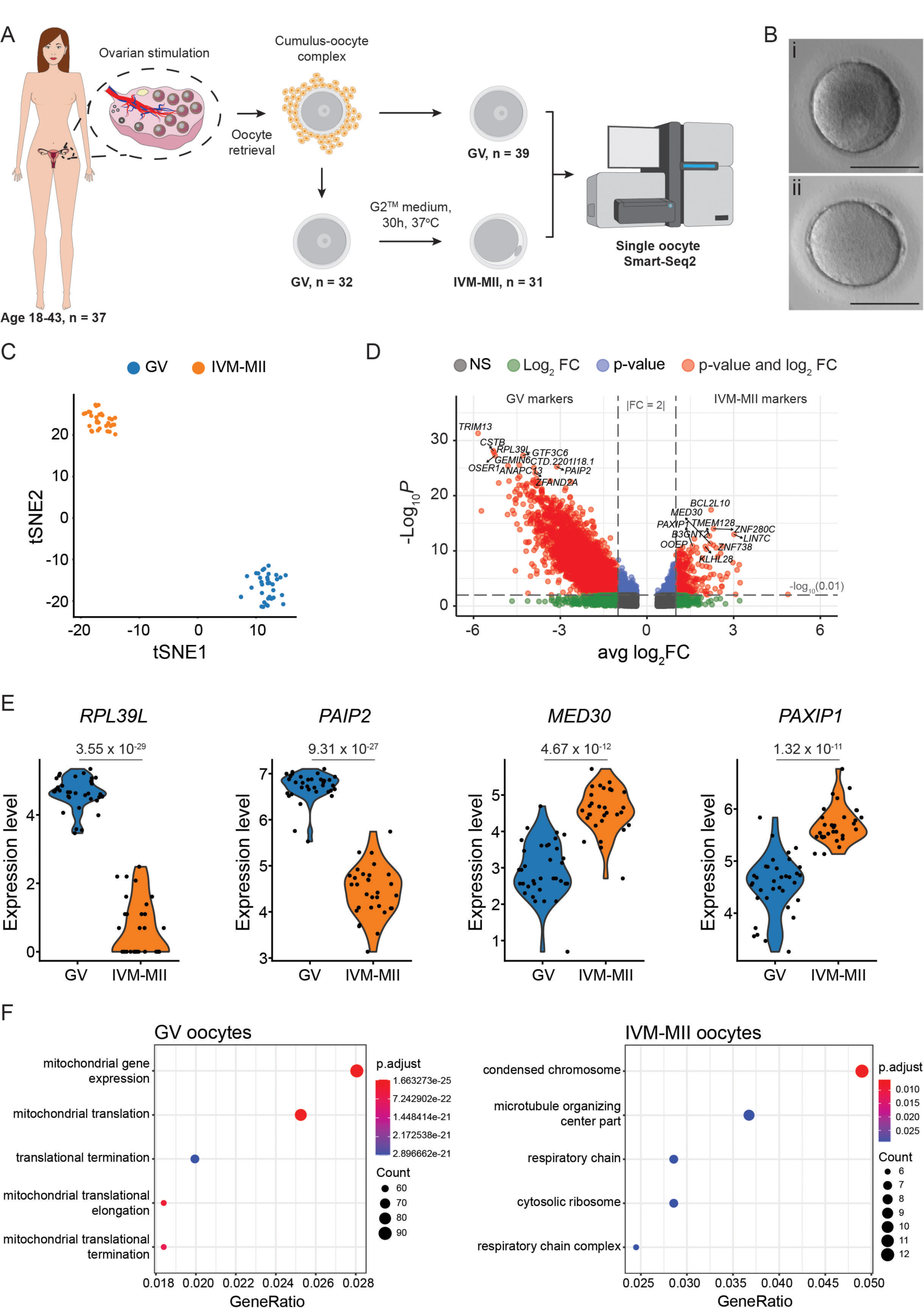
Single-cell transcriptome profiling of human oocytes. **(A)** Schematic representation of the experimental design. Briefly, 37 women were recruited. The mean woman age was 28.8 years (SD = 7.7, range 18 - 43), and from each woman we included between 1-4 GV oocytes. GV oocytes were analysed directly as GV (n=40) right after denudation or, after 30h in G2^™^ medium, as *in vitro* matured metaphase II (IVM-MII, n=32) oocytes. Their transcriptome was compared by single cell RNA-sequencing analysis (Smart-seq2). **(B)** Exemplary pictures of a germinal vesicle (i) and a IVM-MII (ii) oocyte included in the study. Scale bar = 100µm **(C)** Oocytes cluster according to their maturation stage. **(D)** Differentially expressed genes between the GV and the IVM-MII groups are represented in red. Labels correspond to the top10 differentially expressed genes in each category after filtering for fold change 2 (avg_log2FC > 1) and sorting markers according to their p-value (cutoff = 0.01). Total number of variables: 12,431. **(E)** Example of two GV markers (*RPL39L*: Ribosomal Protein L39-Like; *PAIP2*: Poly(A) binding protein Interacting Protein 2) and two IVM-MII markers (*MED30*: MEDiator complex subunit 30; *PAXIP1*: PAX Interacting Protein 1). **(F)** Gene Ontology enrichment analysis of each maturation stage. The top 5 activated GO terms are shown. p-values were adjusted using the FDR method. GV: Germinal Vesicle; IVM-MII: *in vitro* matured metaphase II.

### Oocytes cluster according to maturation stage

First we wanted to know which parameter had the biggest impact on the oocyte transcriptome when performing unbiased clustering of our single oocyte expression data. Dimension reduction analysis by t-SNE revealed two groups of oocytes, with maturation stage being the differentiating feature between clusters (Fig. 1C). After filtering for fold change (FC) > |2| and p-value < 0.01, we identified differentially expressed genes (DEGs) as cluster markers corresponding to each of the maturation stages. Thereby we identified 4,445 GV markers (out of 11,603 ± 373 SEM detected genes in GV oocytes) and 324 IVM-MII markers (out of 8,586 ± 435 SEM detected genes in IVM-MII oocytes) (Fig. 1D, E, Supplementary Table 2, Supplementary Fig. S3). The top 10 genes identified as GV markers ranked by p-value were *TRIM13* (Tripartite Motif Containing 13), *CSTB* (Cystatin B), *RPL39L* (Ribosomal Protein L39 Like), *GTF3C6* (General Transcription Factor IIIC Subunit 6), *OSER1* (Oxidative Stress Responsive Serine Rich 1), *PAIP2* (Poly(A) Binding Protein Interacting Protein 2), *GEMIN6* (Gem Nuclear Organelle Associated Protein 6), *ANAPC13* (Anaphase Promoting Complex Subunit 13), *CTD*.*2201I18*.*1* and *ZFAND2A* (Zing finger AN1-Type Containing 2A). For the IVM-MII oocytes, the top10 markers identified are *BCL2L10* (BCL2 Like 10, apoptosis regulator), *ZNF280C* (Zinc Finger Protein 280C), *LIN7C* (Lin-7 Homolog C, Crumbs Cell Polarity Complex Component), *TMEM128* (Transmembrane Protein 128), *B3GNT2* (UDP-GlcNAc:BetaGal Beta-1,3-N-Acetylglucosaminyltransferase 2), *MED30* (Mediator Complex Subunit 30), *OOEP* (Oocyte Expressed Protein), *PAXIP1* (PAX Interacting Protein 1), *ZNF738* (Zinc Finger Protein 738) and *KLHL28* (Kelch Like Family Member 28).

We then performed gene ontology (GO) term enrichment analysis on the identified maturation stage markers to compare the transcriptomes between GV stage and IVM-MII stage (Fig. 1F). GV oocytes showed a higher abundance in transcripts of genes enriched in GO-terms related to mitochondrial gene expression (e.g. many genes constituting the large and small mitochondrial ribosomes such as *MRPL27* and *MRPS22*). On the other hand, IVM-MII oocytes had more abundant transcripts enriched in GO-terms related to chromosome condensation (e.g. CENPK, ADD3) and microtubule organising center (e.g. CETN3, KIF3A) (Supplementary Table 3).

In summary, unguided clustering of our data identified maturation stage as the main differentiator between oocytes. We identified DEGs for GV and IVM-MII oocytes and revealed GO pathways enriched for each stage.

### Transcript levels change with age for specific gene groups

With maturation stage being the main variable distinguishing our two cell clusters (Fig. 1C), we decided to perform dimension reduction analysis by t-SNE on each one (GV and IVM-MII) separately to determine whether age (i.e. young vs. old women) could be a differentiating feature within each maturation stage. However, this analysis did not reveal separate age-related clusters within GV oocytes or IVM-MII oocytes (Fig. 2A). Furthermore, neither the number of RNA-molecules nor of expressed genes detected per oocyte significantly changed with age (Supplementary Fig. S2), providing additional evidence that the oocyte transcriptomes did not change globally with age.

**Figure 2.**
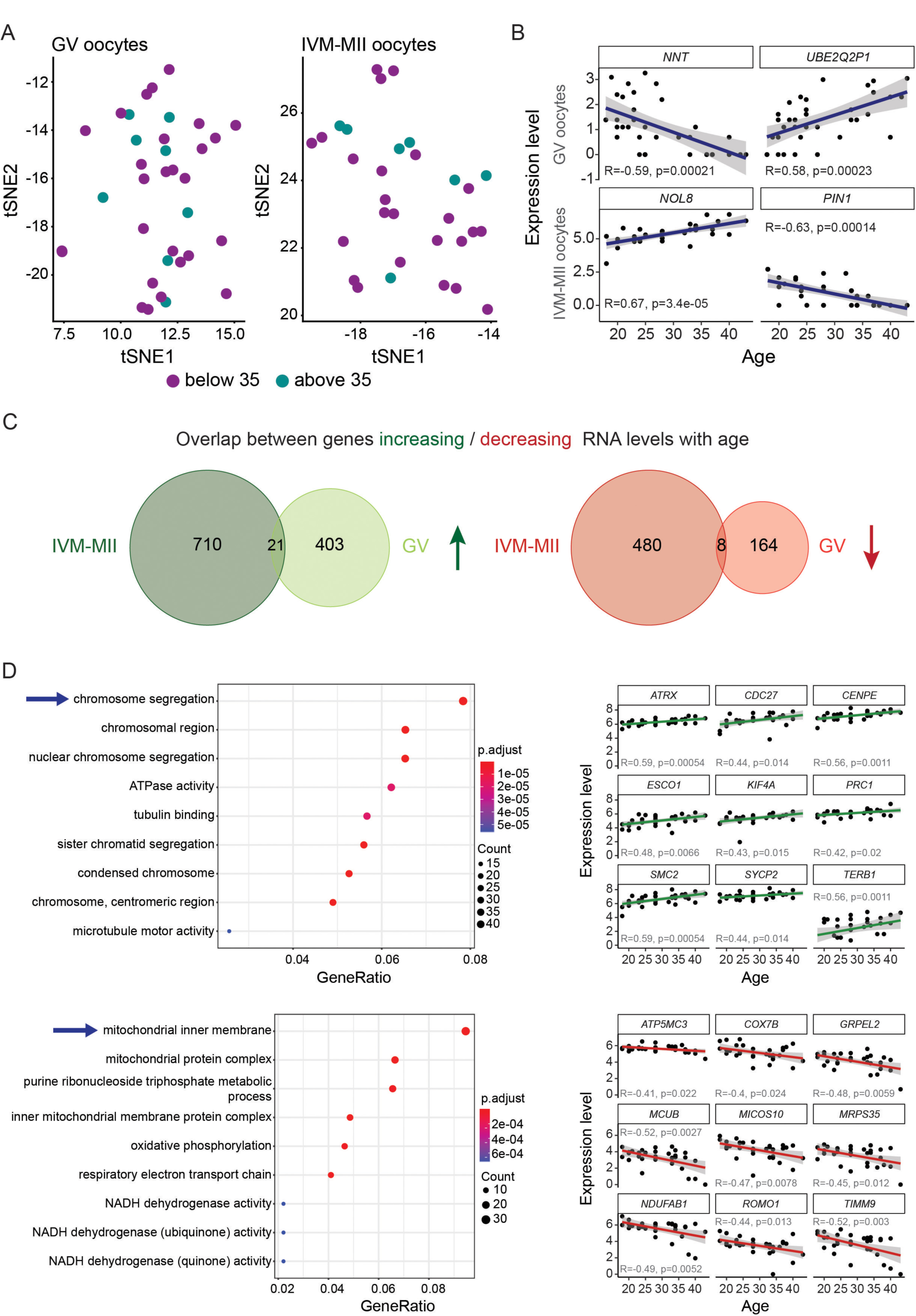
Analysis of gene expression correlation with age. **(A)** Cluster analysis of oocytes at each maturation stage, GV and IVM-MII, independently. **(B)** Examples of genes that correlate with age. *NNT* and *UBE2Q2P1* are genes found in the GV set of oocytes, while *NOL8* and *PIN1* were identified within the IVM-MII oocyte population. **(C)** Venn diagrams showing the overlap between genes changing RNA levels with age in GV and IVM-MII oocytes. In green, genes whose RNAs abundance increased with age, in red the number of genes whose RNAs abundance decreased with age. **(D)** Gene ontology analysis of genes that change with age in IVM-MII oocytes (left) and examples of how expression levels of genes in some of the most significant GO terms (indicated by arrows) vary with age (right). Upper panel: genes belonging to the GO term “chromosome segregation” for which expression increases with age. Lower panel: genes belonging to the GO term “mitochondrial inner membrane” for which expression decreases with age.

Next, as oocyte quality declines with age, we sought to identify specific genes, of which RNA-levels would increase or decrease in an age-dependent manner. For that reason, we performed correlation tests between gene expression and age independently for GV and IVM-MII oocytes. For each detected gene we obtained a Pearson correlation value (R) and a p-value. The genes that presented an absolute correlation value equal or over 0.3 (R => |0.3|) and a p-value lower than 0.05 were considered as genes for which transcript levels correlated either positively (increased) or negatively (decreased) with age (Supplementary Table 4). Following these criteria, we identified a total of 596 genes changing with age within the GV population and 1,219 genes within the IVM-MII population. For GV oocytes, the top 5 genes (according to R value) that increase in RNA levels with age are *ENSG00000278292* (Ret Finger Protein Like 4A Pseudogene 6), *UBE2Q2P1* (Ubiquitin Conjugating Enzyme E2 Q2 Pseudogene 1), *WFIKKN2* (WAP, Follistatin/Kazal, Immunoglobulin, Kunitz And Netrin Domain Containing 2), *LINC02022* (Long Intergenic Non-Protein Coding RNA 2022) and *ENSG00000227240* (lncRNA, new transcript) and the top 5 genes decreasing the amount of transcript with age are *NNT* (Nicotinamide Nucleotide Transhydrogenase), *DCK* (Deoxycytidine Kinase), *KAT8* (Lysine Acetyltransferase 8), *RG518* and *ARHGEF26* (Rho Guanine Nucleotide Exchange Factor 26) (Fig. 2B). For the IVM-MII oocytes, the top 5 genes that increase / decrease the amount of RNA levels with age are *NOL8* (Nucleolar Protein 8), *TNIK* (TRAF2 And NCK Interacting Kinase), *ESCO2* (Establishment Of Sister Chromatid Cohesion N-Acetyltransferase 2), *AFDN* (Afadin, Adherens Junction Formation Factor), *TPR* (Translocated Promoter Region, Nuclear Basket Protein) and *PIN1* (Peptidylprolyl Cis/Trans Isomerase, NIMA-Interacting 1), *SERF2* (Small EDRK-Rich Factor 2), *C12orf75* (Chromosome 12 Open Reading Frame 75), *TLR5* (Toll Like Receptor 5), *KIAA1671* (Uncharacterized Protein KIAA1671) respectively.

Then we created Venn diagrams, in order to see to which degree the genes whose RNA levels positively or negatively correlated with age overlapped between GV and IVM-MII oocytes (Fig. 2C). To our surprise, we found very little overlap between the 424 genes in GV oocytes and the 731 genes in IVM-MII oocytes whose RNA levels increased with age, with only 21 genes increasing both in GV and IVM-MII oocytes. On the other hand, 488 and 172 genes decreased RNA levels in IVM-MII and GV with age, respectively, with only 8 genes commonly decreasing their RNA levels in both. This suggests that age affects the RNA-levels of different genes in GV and IVM-MII oocytes.

After this preliminary analysis, the genes present in the correlation lists were further analysed using clusterProfiler (Yu *et al*., 2012). We looked for enriched GO terms in the lists of genes that increased in RNA level with age separately from those whose RNA levels decreased with age. When analysing the GV stage oocytes we did not find any particular GO term significantly enriched (p<0.05) after adjusting the p-values (FDR method), neither in the set of genes increasing RNA levels with age nor in the ones decreasing RNA levels with age. In contrast, in the GO analysis performed on genes whose transcript levels changed with age on the IVM-MII stage oocytes we did find significantly enriched GO terms after adjusting the p-value (FDR method) (p < 0.05) (Fig. 2D, left panels). The most enriched GO terms we found within the genes which increased in transcript levels with age were predominantly related to chromosome segregation whereas the most enriched GO terms for the set of genes presenting lower transcript levels with age were mostly related to mitochondrial function (Supplementary Table 5). Examples of specific genes belonging to the most significantly enriched GO term, “chromosome segregation” for genes positively correlating (RNA levels increase) with age were *ATRX* (ATRX Chromatin Remodeler), *CDC27* (Cell Division Cycle 27), *CENPE* (Centromere Protein E), *ESCO1* (Establishment Of Sister Chromatid Cohesion N-Acetyltransferase 1), *KIF4A* (Kinesin Family Member 4A), *PRC1* (Protein Regulator Of Cytokinesis 1), *SMC2* (Structural Maintenance Of Chromosomes 2), *SYCP2* (Synaptonemal Complex Protein 2) and *TERB1* (Telomere Repeat Binding Bouquet Formation Protein 1). On the other hand, specific genes that negatively correlated (RNA levels decrease) with age and that belonged to the “mitochondrial inner membrane” GO term were *ATP5MC3* (ATP Synthase Membrane Subunit C Locus 3), *COX7B* (Cytochrome C Oxidase Subunit 7B), *GRPEL2* (GrpE Like 2, Mitochondrial), *MCUB* (Mitochondrial Calcium Uniporter Dominant Negative Subunit Beta), *MICOS10* (Mitochondrial Contact Site And Cristae Organizing System Subunit 10), *MRPS35* (Mitochondrial Ribosomal Protein S35), *NDUFAB1* (NADH:Ubiquinone Oxidoreductase Subunit AB1), *ROMO1* (Reactive Oxygen Species Modulator 1) and *TIMM9* (Translocase Of Inner Mitochondrial Membrane 9) (Fig. 2D, right panels).

In order to assess to which degree maturation stage markers for GV or IVM-MII oocytes also changed in RNA abundance with age, we intersected stage and age markers through Venn diagrams (Supplementary Fig. S4). We observed that only a small proportion of maturation stage markers changed with age and that the majority of transcripts changing with age were not from maturation stage markers (only 22% for GV and 3% for IVM-MII). This suggests that maturation stage identity (GV vs. IVM-MII), in line with its importance for reproduction, is largely preserved during oocyte ageing. In summary, we found that RNA levels of different genes changed with age in GV and IVM-MII oocytes. In particular in IVM-MII oocytes, we found transcripts related to chromosome segregation and mitochondrial function to be affected by age, both pathways, which have been implicated in the age-related oocyte quality decline (Nagaoka *et al*., 2012; May-Panloup *et al*., 2016).

### Upstream master regulators differ between up- and downregulated genes

Considering that the IVM-MII stage is where we found more changes in RNA abundance correlated with age and significantly enriched GO terms in genes both increasing and decreasing tendency, we performed a gene regulatory network analysis using the R package Genie3 (Huynh-Thu *et al*., 2010). As input we gave an expression matrix where only IVM-MII stage oocytes and the genes whose RNA levels correlated (increased/decreased) with age within this maturation stage were included. Moreover, we used a list of described human transcription factors (Lambert *et al*., 2018) as potential regulators of the network. When plotting the results of this analysis (Fig. 3A, Supplementary Table 6) we took into account the results obtained from the previous GO analysis (Fig. 2D). From this analysis it became clear that the upstream potential master regulators for each of the GO terms were different. We observed GC-Rich Promoter Binding Protein 1 (*GPBP1*) and RLF zing finger (*RLF*) as potential regulators of genes related to “chromosome segregation” while basonuclin 1 (*BNC1*), thyroid hormone receptor beta (*THRB*) and transcription termination factor 1 (*TTF1*) appeared to be mostly regulating genes related to “mitochondrial inner membrane”. Moreover, SON DNA and RNA binding protein (*SON*), which is a splicing factor belonging to the “RNA splicing” GO term, appears as well as one of the main regulators of the network. The expression dynamics of the mentioned potential master regulators mostly follows the tendency of the genes within the GO term they regulate. For example, *GBPB1* and *RLF*, which we identified as potential upstream regulators of the GO term “chromosome segregation”, increased in RNA levels with age, as do the genes present in this particular GO term (Fig. 3B). In the case of the potential upstream master regulators of the GO term “mitochondrial inner membrane”, identified within the genes decreasing in transcript levels with age, *BNC1* and *THRB* followed the same dynamics, while *TTF1* behaved the opposite, with its transcript levels increasing with age (Fig. 3B).

**Figure 3.**
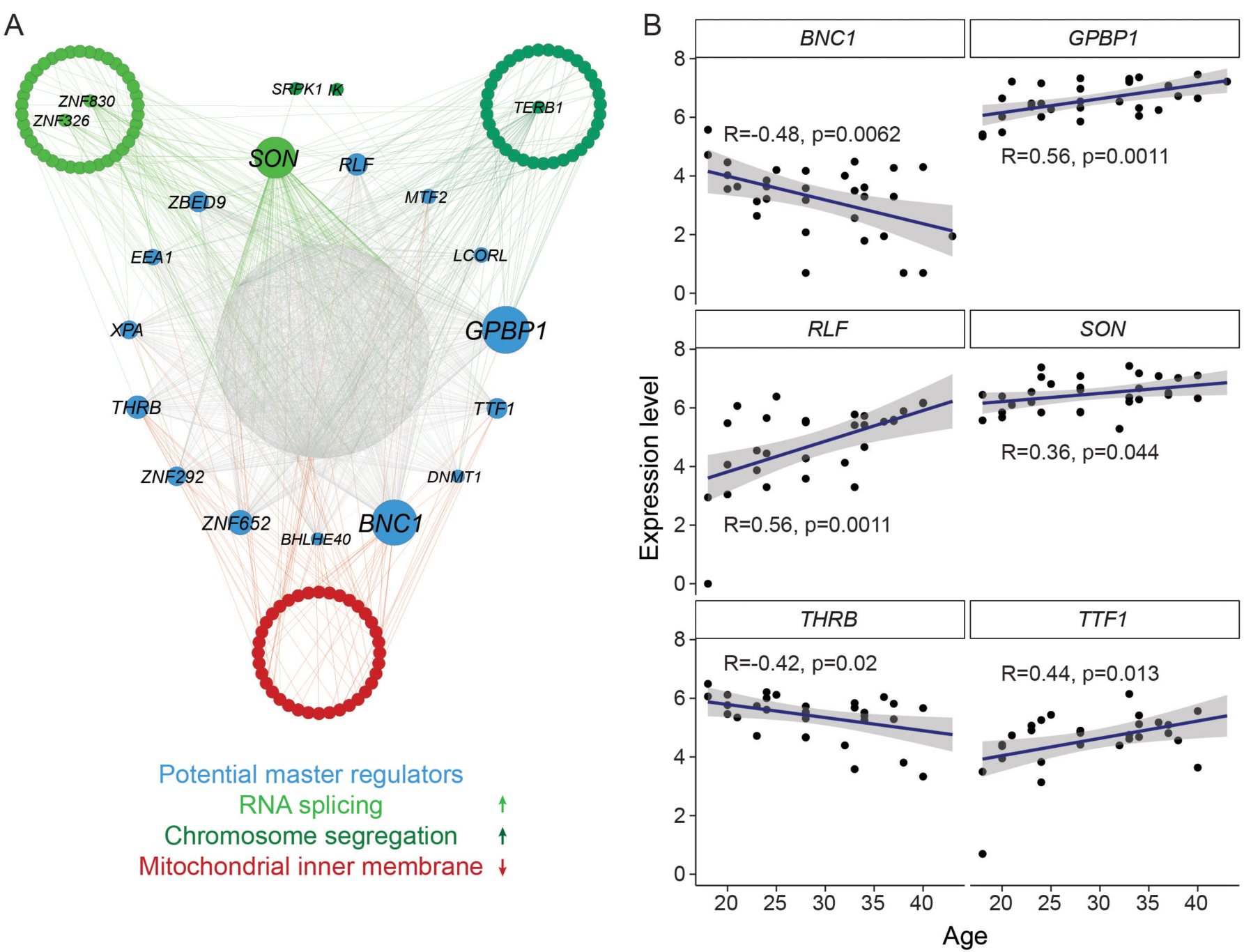
Gene regulatory network analysis. **(A)** Cytoscape plots from the top 2,500 regulatory links among genes found to correlate with age in IVM-MII oocytes. A list of known human transcription factors was given as an input to Genie3 to be used as potential regulators of the network. In green are genes belonging to two of the GO terms enriched in genes whose RNA levels increase with age, “RNA splicing” (light green) and “chromosome segregation” (dark green). *SRPK1* and *IK* belong to both of these GO terms, and therefore they are plotted in between. In red are genes for which RNA levels decrease with age belonging to the enriched GO term “mitochondrial inner membrane”. **(B)** Expression dynamics of the potential master regulators of genes that correlate with age in IVM-MII oocytes.

Taking into account that not only transcription factors but also other groups of proteins such as those involved in RNA stability or RNA processing might play a role in regulation of gene expression and RNA abundance, we decided to perform the Genie3 analysis also in an unbiased manner. Instead of giving as an input a predetermined list of potential regulators of defined categories like transcription factors, we instead allowed all genes of the network to be potential regulators (Supplementary Fig. S5A, Supplementary Table 6). Interestingly, the main nodes that we obtained in this analysis mostly differed from the ones described above, except from *SON* and *RLF*, which appeared again as main nodes in the regulatory network. Among the main regulators that we found in this analysis were genes involved in transcriptional regulation (*CLOCK, DHX9, ZKSCAN5, UHRF1*), genes related to RNA regulation (*SON, EDC3)*, genes related to DNA damage response (*PDCD5*) and genes that regulate centrosome and mitotic spindle integrity (*HAUS7*) or that might influence mitochondrial activity (*VDAC3*). The expression dynamics of the genes in relation to age is shown in Supplementary Fig. S5B.

Taken together, we hereby identified a number of potential master regulators, which could determine the observed RNA level changes of specific groups of genes in relation to age in IVM-MII oocytes.

### Impact of Body mass index (BMI) on human oocyte transcriptome

Both obesity (Shah *et al*., 2011; Machtinger *et al*., 2012) as well as underweight (Brower *et al*., 2013) in women have been associated with poor oocyte quality and reproductive outcome. Taking advantage of the availability of BMI information from each woman included in our study, we analysed whether this factor could also influence human oocyte quality at the transcriptome level. As we did for age, we considered BMI as a continuous variable and looked at the correlation between BMI and AFC as well as BMI and gene expression. Our dataset did not show any correlation between BMI and AFC (Fig. 4A; R = 0.053, p-value = 0.64).

**Figure 4.**
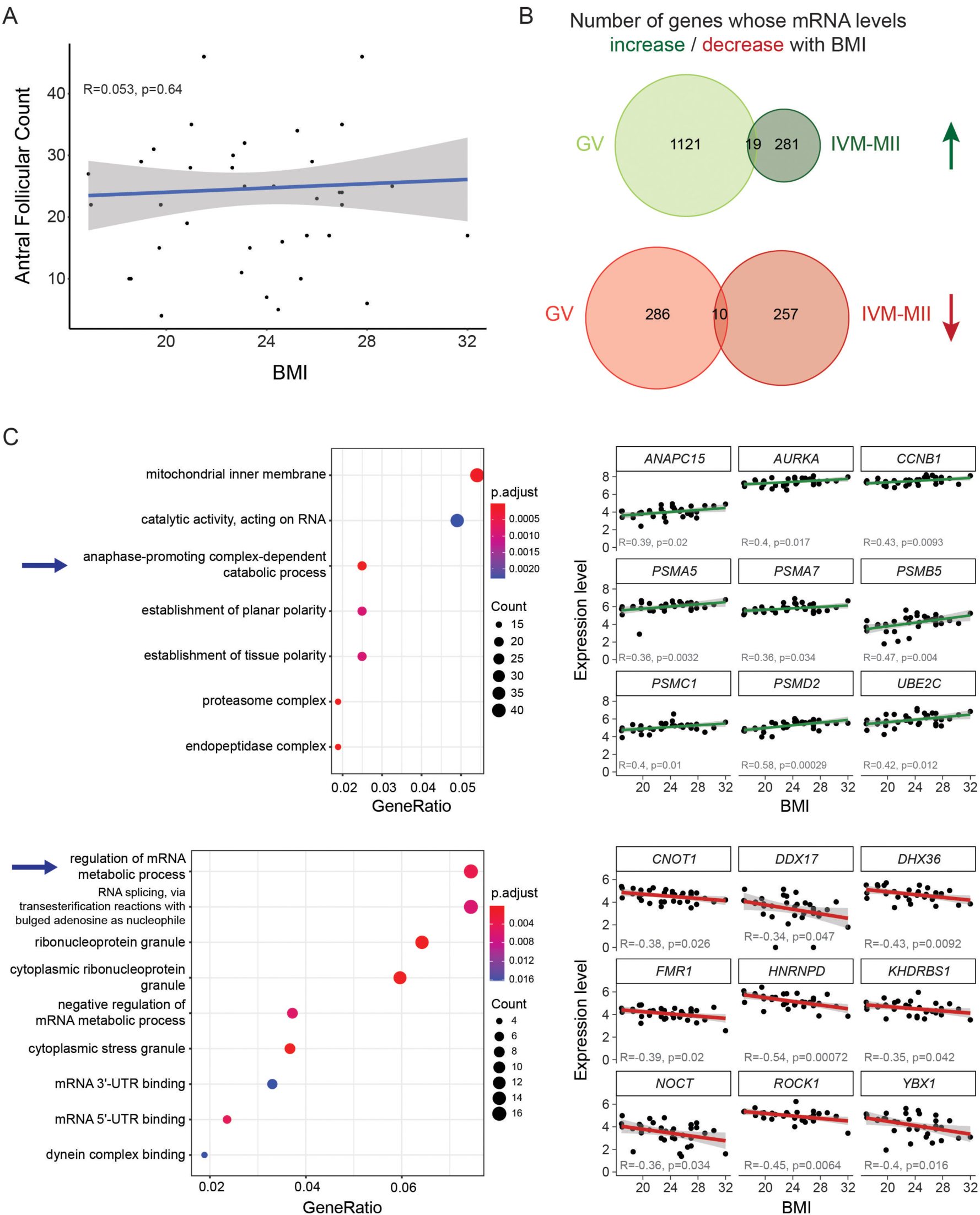
Analysis of gene expression correlation with BMI. **(A)** Correlation plot between AFC and BMI. **(B)** Venn diagrams showing the number of genes whose amount of RNA correlates with BMI in GV and IVM-MII oocytes. In green, genes whose RNA abundance increases with BMI, in red the number of genes whose RNA abundance decreases with BMI. **(C)** Gene ontology analysis of genes that correlate with BMI in GV oocytes (left) and examples of how expression levels of genes in some of the most significant GO terms (demarcated by arrow) vary with age (right). Upper panel: genes for which expression increases with BMI. Lower panel: genes for which expression decreases with BMI.

In terms of gene expression, again we analysed GV- and IVM-MII-stage oocytes independently. As opposed to age, BMI influenced the transcript levels of GV oocytes more than it did for the IVM-MII oocytes. In the GV oocytes we found a total of 1,436 genes whose transcript levels correlated with BMI, mostly (79.4%) positively (Fig. 4B, Supplementary Table 7). In the case of IVM-MII oocytes, a total of 567 genes were found to change in transcript levels with BMI. In this case, about half of the genes (300) positively correlated with BMI while the other half (267) negatively correlated with BMI. As it was the case for RNA-level changes associated with age (Fig. 2C), also in this case, we observed almost no overlap between genes whose RNA-levels changed in GV or IVM-MII oocytes in relation to BMI (Fig. 4B).

Gene ontology analysis revealed no specific term enriched within the list of genes correlating with BMI in the IVM-MII oocytes. For GV oocytes, gene ontology analysis revealed that the list of genes increasing transcript levels with increasing BMI is enriched in GO terms like “mitochondrial inner membrane”, “catalytic activity, acting on RNA”, “anaphase-promoting complex-dependent catabolic process” and “establishment of (planar/tissue) polarity” among others (Fig. 4C, Supplementary Table 8). For genes that decreased in RNA level with rising BMI in GVs, gene ontology analysis revealed terms mostly related to regulation of (m)RNA metabolism and RNA splicing.

Overall, BMI affected oocyte transcript levels of specific pathways especially in GV oocytes, which potentially would contribute to the decline in oocyte quality and fertility in women with abnormal BMI.

## Discussion

### This study in light of previous work

In this study we have investigated the effect of age on oocyte quality at the RNA level. We have used Smart-seq2 (Picelli et al. 2013) single-cell RNA-Sequencing technology instead of microarrays or bulk RNA-seq protocols commonly used in previous studies. Moreover, while comparable recent studies on the impact of oocyte age on the human oocyte transcriptome analyzed only a small number of oocytes (Hendrickson *et al*., 2017; Reyes *et al*., 2017; Zhang *et al*., 2020), our dataset includes a large number of single oocytes (n=72) from a large cohort of women of a wide-ranging age-span (n=37, 18-43 years), thereby increasing our statistical power in comparison to those previous studies.

The design of our study allowed us to compare the transcriptomes of single GV and IVM-MII oocytes obtained from women of different ages. Besides the identification of maturation stage-specific markers, our dataset allowed us to analyse oocyte age-related quality decline by correlation analysis over time rather than comparing oocytes above or below an arbitrary age threshold as done in previous studies. In addition, our dataset allowed us to identify RNAs correlating with BMI.

### Maturation stage is the main differentiator of oocyte transcriptomes

Despite the identification of transcriptome changes associated with reproductive ageing, we show that age is not the main driver of transcriptomic variability in oocytes. Instead, maturation stage (GV vs. IVM-MII) was the main factor clearly separating our pool of oocytes into two distinct populations in a tSNE cluster analysis (Fig. 1C, 2A). Looking at the differentially expressed genes between these two maturation stages, we identified a higher number of genes presenting increased RNA abundance within the GV population (4,445) in comparison to IVM-MII oocytes (324), which is not surprising considering that during nuclear maturation from fully grown GV to IVM-MII stage there is no (or very little) transcription (De La Fuente, 2018). From the stage markers, which we have identified in our dataset at the RNA-level, some have been previously also found to be stage-specific GV (TDRKH) and MII (WEE2, DNMT1) protein markers by single-cell proteomics (Supplementary Fig. S3) (Virant-Klun *et al*., 2016). This confirms that the differences we observed in RNA-abundance are also reflected at the protein level at least for these selected marker genes.

In mouse oocytes, Su and colleagues found that during maturation from GV to MII stage, transcripts were degraded on a large scale and only those transcripts related to pathways essential for the maintenance of the unique MII characteristics were protected selectively from degradation (Su *et al*., 2007). On the other hand, transcripts responsible for meiotic arrest at GV stage and progression of oocyte maturation were degraded in MII oocytes in mice. Those included transcripts related to oxidative phosphorylation, energy production, protein synthesis and metabolism. We made similar observations in human oocytes. In our dataset we observed terms related to protein synthesis and mitochondria enriched at the GV stage. In consequence, and considering that we compared immature (GVs) vs. *in vitro* matured (IVM-MII) oocytes, our results lead us to infer a suppression of translation-related terms within the IVM-MII oocyte population, indicating a shut down of protein synthesis during maturation, as has been described in mouse oocytes (Su *et al*., 2007). Furthermore, we observed enrichment of genes related to chromosome condensation in IVM-MII oocytes, in line with the establishment of metaphase chromosomes at this maturation stage. Therefore, our data support the selective degradation of GV-specific RNA-molecules and protection from degradation of MII-specific transcripts previously observed in mice during oocyte maturation, indicating that this process is conserved during human oocyte maturation from GV to IVM-MII stage.

### Transcripts affected by age are related to oxidative stress, mitochondrial function, chromosome segregation and DNA metabolism pathways

In our dataset, over 1,700 genes changed in transcript levels in correlation with women’s age. In general, the majority of transcripts changing with age were not maturation stage markers supporting the idea that ageing does not affect overall cell identity (Wang *et al*., 2020). Instead, age has an impact on the transcript levels of specific groups of genes with importance for oocyte function, explaining the decline in oocyte quality with age (Wang *et al*., 2020).

We observed that the transcriptome of IVM-MII stage oocytes was more affected by age than those of GV oocytes, based on the number of genes with increased or decreased RNA levels. This trend has been also previously observed in mouse oocytes (Pan *et al*., 2008). Furthermore, we also observed that the genes whose expression levels correlated with age differed between GV and IVM-MII stage, which is in agreement with a recent study on non-human primate oocytes (Wang *et al*., 2020) where genes differentially expressed with age were found to be oocyte maturation stage-specific. Among the few genes whose RNA levels increased with age in both GV and IVM-MII oocytes, we found *FAM210B*, a mitochondrial factor that has been associated with human ovarian cancer (Sun *et al*., 2017). Further examples were *TERB1*, which encodes a meiosis-specific telomere-associated protein involved in attaching the meiotic telomere to the inner nuclear membrane, and *RFC1*, encoding a subunit of the replication factor C, a DNA polymerase accessory protein required for DNA replication and repair and that might also play a role in telomere stability. Interestingly, none of these three genes varied significantly in RNA-level between GV and IVM-MII stage, suggesting that their age-related increase in RNA levels was already present at the GV stage and was maintained through the *in vitro* maturation step to the IVM-MII stage. Among the genes whose RNA levels decreased with age in both GV and IVM-MII stages, we found *ND1*, which encodes the mitochondrial NADH:Ubiquinone Oxidoreductase Core Subunit 1 and is involved in electron transport in the respiratory chain.

Even though we did not find any specific GO term enriched within the group of genes correlating with age in GV oocytes, we did find levels of the oxidative stress-related *GPX1* RNA to decrease with age, corresponding with what has been recently shown for non-human primate oocytes (Wang *et al*., 2020). Furthermore, we observed *RPA1* RNA levels increasing with age. This gene plays an important role in DNA metabolism including replication, repair and recombination among other functions. Moreover *RPA1* has been previously related to ageing (present in the GenAge database, https://genomics.senescence.info/, (Tacutu *et al*., 2018)). Other examples of genes related with ageing according to the GenAge database and that we also found in our set of IVM-MII oocytes were the topoisomerase gene *TOP2B* and the antioxidase *PRDX1*. Surprisingly, *TOP2B* has been previously found to decrease in human oocytes with age (Zhang *et al*., 2020), while we observed the opposite trend. A potential origin for such a discrepancy is that Zhang and colleagues studied only 3 *in vivo* MII oocytes for young (<30 years old) women and 3 for older (>40 years old) women. On the other hand, we used a total of 31 IVM-MII oocytes from women ranging from 18 to 43 years of age, and instead of comparing arbitrary age groups performed a correlation analysis of transcript levels with age. However, we are aware of the limitations that our study has when using *in vitro* matured oocytes. *PRDX1*, which belongs to the peroxiredoxin family of antioxidant genes has been suggested to play an antioxidant protective role. Interestingly, we found genes related to oxidative stress protection decrease with age, in both GV oocytes (*GPX1*) and IVM-MII oocytes (*PRDX1*). Indeed, it is known that with age, there is an increase of oxidative molecules and a decrease in the expression of oxidation protective genes. This phenomenon is known as the “oxidative stress theory of ageing” (Liguori *et al*., 2018). A decrease in the expression of genes responsible for protecting against oxidative stress has been previously observed as a general feature of ageing mouse, non-human primate and human oocytes (Lim and Luderer, 2011; Reyes *et al*., 2017; Wang *et al*., 2020).

The gene ontology analysis we performed on the list of genes whose transcript levels correlate with age in the IVM-MII oocytes revealed that the terms significantly enriched among the genes which alter their transcript levels with age could be classified into the following: chromosome segregation, cell cycle regulation, mitochondrial function and RNA metabolism. All of these biological processes have been previously reported to be altered with age in the human oocyte (Steuerwald *et al*., 2007; Grøndahl *et al*., 2010; Almansa-Ordonez *et al*., 2020). In fact, if we take a closer look at the entire list of GO terms enriched in the set of genes either with increased or decreased RNA levels with age, we detect both, discrepancies and correlations with what has been previously published regarding the relationship between the human oocyte transcriptome and female age (Steuerwald *et al*., 2007; Grøndahl *et al*., 2010). For example, (Grøndahl *et al*., 2010) found “DNA repair” to be enriched in old MII human oocytes, and we also found “regulation of DNA repair” among the GO terms enriched in genes increasing their RNA levels with age. Another example in which our data correlates with what has been published before is the GO term “mitochondrial membrane”, which was also found to be downregulated in older oocytes (Steuerwald *et al*., 2007). Nonetheless, in this same publication they found the GO term “cell cycle checkpoint” to be downregulated in older oocytes, while we find it in the set of genes increasing their RNA level with age.

Besides the GO terms specifically enriched in the lists of genes whose transcript levels vary with age, we also found individual genes known to be involved in oocyte maturation that decreased in transcript levels with age, which might be another indication of low quality oocytes. Examples of such genes are the growth differentiation factor 9 (*GDF9)* and the cytoplasmic polyadenylation element binding protein (*CPEB2*) (Supplementary Table 4). GDF9 is an oocyte-derived paracrine factor that plays a role in the communication between the cumulus cells and the oocyte and is responsible to keep the integrity of the cumulus-oocyte complex in the preovulatory follicles and after ovulation (Yan *et al*., 2001). Even though we don’t have cumulus cells in our *in vitro* maturation system, the fact that we observe a decrease of *GDF9* transcript with age suggests it to be a potential factor in age-related impairment of cumulus-oocyte communication. CPEB2, on the other hand, is a potential regulator of cytoplasmic RNA polyadenylation during oogenesis and has been described to be important for porcine oocyte meiotic maturation to MII stage and early embryogenesis (Prochazkova *et al*., 2018). Cytoplasmic polyadenylation is a mechanism known to regulate stability and translation of maternal-effect mRNAs for protein production during oogenesis (Susor and Kubelka, 2017). Considering that we observe a decrease of *CPEB2* transcript amount with age in IVM-MII oocytes, this might suggest an impairment of old MII oocytes to produce a mature proteome.

Advanced maternal age has been shown to be directly related to an increased rate of aneuploidies, with cohesin loss and centromeric abnormalities being the most studied underlying causes (Tachibana-Konwalski *et al*., 2010; Burkhardt *et al*., 2016; Smoak *et al*., 2016; Toth and Jessberger, 2016; Gruhn *et al*., 2019; Zielinska *et al*., 2019). Strikingly, in our data, among the top hits of biological processes affected by maternal age in IVM-MII oocytes are chromosome and chromatid segregation-related terms (Supplementary Table 5). To name some key examples, the cohesin loader NIPBL and the cohesin release factor WAPL, the cohesin complex member SMC3, structural maintenance of chromosome factors SMC4 and SMC5, the synaptenomal complex member SYCP2 and centromeric proteins like CENPC, CENPE, CENPF, CENPM and INCENP are all increased at the RNA level with age in our IVM-MII oocytes (Supplementary Table 4, Fig. 2D). Specifically, SMC5 for example has been shown to be important for correct segregation of homologous chromosomes during meiosis I in mouse oocytes and, in contrast to our human RNA data, that protein levels of SMC5/6 declined in mice oocytes with age (Hwang *et al*., 2017). Why did we observe an increase in the RNA-levels of chromosome segregation genes with age? A potential explanation could be that ageing oocytes somehow sense the decline of key chromosomal components with age and attempt to compensate for it at the RNA-level. Our results therefore point towards the hypothesis that not only protein stability of key chromosomal factors but also abnormal changes at the RNA level of those genes might contribute to the increased aneuploidy rate with advanced maternal age. Indeed, Fragouli and colleagues described a link between transcriptomic alterations of genes involved in biological processes such as spindle assembly or chromosome alignment and aneuploidy of oocytes (Fragouli *et al*., 2010).

All together, we identified genes linked to oxidative stress decreasing in transcript levels in both, GVs and IVM-MIIs, as well as genes related to DNA metabolism increasing in transcript level with age. Moreover, in IVM-MII oocytes we found genes involved in chromosome segregation or chromatin organization that were upregulated with age, while genes related to mitochondrial function decreased in transcript levels with age.

### Identification of potential master regulators of age-related changes in oocyte transcriptome by network analysis

In order to identify master regulators of the pathways affected by age, we used a network analysis approach (Huynh-Thu *et al*., 2010) using a list of human transcription factors (Lambert *et al*., 2018) as potential regulators of the network. Among the potential upstream regulators in IVM-MII-stage oocytes we identified the zinc finger transcription factor *BNC1*. It is expressed in germ cells of both ovaries and testes and plays a role in the regulation of rRNA transcription (Zhang *et al*., 2007). Specifically, *BNC1* is expressed in oocytes present within secondary follicles and in ovulated oocytes and its deficiency has been associated with premature ovarian failure (Zhang *et al*., 2018). Furthermore, knockdown of BNC1 in human oocytes leads to impaired meiotic maturation and a decrease of the oocyte-derived proteins BMP15 and p-AKT (Zhang *et al*., 2018). In mouse oocytes, Bnc1 knock-down leads to impaired RNA polymerase I and II transcription, altered oocyte morphology and a failure of embryos to develop beyond the 2-cell stage resulting in female subfertility (Ma *et al*., 2006). Finally, Bnc1 deficiency causes premature testicular ageing in mice (Li *et al*., 2020), indicating a general link to germ cell ageing pathways.

Another potential master regulator that we identified in our gene regulatory network analysis is *SON*. This gene encodes an RNA binding protein, which promotes pre-mRNA splicing, specially of transcripts presenting weak splice sites and transcripts related to cell-cycle and DNA repair (Lu *et al*., 2014). SON has also been found to be important for human embryonic stem cell (hESC) pluripotency by regulating proper splicing of pluripotency and germ cell factors such as *OCT4* and *PRDM14 (**Lu et al*., *2013*). Interestingly, *SON* not only belongs to the GO term “RNA splicing”, which is found among genes with increased transcript levels with age, but was also identified in our network analysis as a potential regulator of RNA splicing related genes. This is in line with extensive autoregulatory cross-talk within the splicing machinery (Saltzman *et al*., 2011; Papasaikas *et al*., 2015).

### Transcriptomic changes associated with BMI

Besides age, as well BMI is a main driver affecting oocyte quality and female reproductive fitness (Shah *et al*., 2011; Machtinger *et al*., 2012; Brower *et al*., 2013). Therefore we made use of the BMI information recorded for each donor in our dataset and correlated BMI with RNA levels. Among the enriched GO terms for RNAs increasing with BMI in GV oocytes, we found “anaphase-promoting complex-dependent catabolic process”, which includes the anaphase promoting complex subunits *ANAPC11* and *ANAPC15*, as well as *AURKA* (Aurora kinase A), which is important for microtubule nucleation during meiotic spindle assembly in mouse oocytes (Saskova *et al*., 2008; Solc *et al*., 2012; Namgoong and Kim, 2018). This could explain previously reported spindle abnormalities in oocytes from obese women (Machtinger *et al*., 2012).

In addition, RNA splicing and other GO terms related to RNA metabolism/dynamics were not only influenced by age in our dataset, but also by BMI in GV stage oocytes, where we observed a decrease in RNA levels for genes associated with these pathways. An example is the downregulation with increasing BMI of CNOT1 (CCR4-NOT Transcription Complex Subunit 1), a scaffolding unit of the CCR4-NOT complex, which is involved in RNA-deadenylation, RNA-degradation and translational repression (Shirai *et al*., 2014). The CCR4-NOT complex has been implicated in the selective deadenylation and degradation of transcripts during meiotic maturation in mouse oocytes and the disruption of the CNOT6L subunit of the CCR4-NOT complex caused defects in microtubule-chromosome organization and resulted in meiotic arrest (Sha *et al*., 2018; Vieux and Clarke, 2018). Another RNA regulator changing with BMI we wanted to highlight is *FMR1* (fragile X mental retardation 1), which is an inhibitor of mRNA translation, and if mutated causes fragile X-associated primary ovarian insufficiency (FXPOI) (Sherman *et al*., 2014). This syndrome is characterized by premature ovarian failure (POF) resulting in infertility or early menopause in women as well as impaired fertility and ovarian abnormalities in mouse models (Hoffman *et al*., 2012; Lu *et al*., 2012).

### Conclusion

In this study we have shown that age as well as BMI affect key pathways at the RNA-level, which are involved in oocyte maturation and function such as chromosome segregation, mitochondria, RNA-metabolism and translation. While many age-related oocyte defects, as for example chromosome segregation errors, have been attributed to low protein turnover during meiotic arrest (Tachibana-Konwalski *et al*., 2010; Burkhardt *et al*., 2016; Smoak *et al*., 2016; Toth and Jessberger, 2016; Gruhn *et al*., 2019; Zielinska *et al*., 2019), it has been less appreciated that as well the transcripts related to centromeric- or cohesin-associated factors are misregulated during oocyte ageing. Maturing oocytes go through a dramatic rewiring of gene expression dynamics, which includes phases of global transcriptional and translational repression and selective RNA polyadenylation and RNA degradation (Clarke, 2018; De La Fuente, 2018). As we found changes in transcript levels for RNA-processing genes both with age and BMI, this might in turn lead to impaired RNA-levels and processing of critical oocyte maturation genes needed at the time. Thereby changes in the transcriptome could contribute to the low oocyte quality in both women with advanced maternal age and abnormal BMI.

In summary, we provide with this study a high resolution analysis of the pathways affected at the RNA-level in correlation with age and BMI in GV and IVM-MII oocytes. Besides advancing our knowledge on the underpinnings of the age- and BMI-related oocyte quality decline, we deliver a rich resource for the field, guiding further investigations and potential diagnostic or therapeutic developments related to oocyte quality.

## Author’s roles

M.B., A.M., H.H., R.V. and B.P. conceived the study. M.B. collected and performed IVM on oocytes. P.L. and S.R. performed single cell sequencing. S.L., P.N. and M.E. performed Bioinformatic analysis. F.Z. assisted with data analysis and revised the manuscript B.P., R.V. and H.H. acquired funding and supervised the project. S.L., M.B. and B.P. wrote the paper with input from all other authors.

## Acknowledgements

We would like to acknowledge all members of the Heyn, Vassena and Payer labs for their input and discussions and Elvan Boke for expert reading of the manuscript.

## Data availability

The data underlying this article are being uploaded to the Gene Expression Omnibus (GEO) and will be made available with publication of the manuscript.

## References

Almansa-Ordonez A, Bellido R, Vassena R, Barragan M, Zambelli F. Oxidative Stress in Reproduction: A Mitochondrial Perspective. Biology [Internet] 2020;9.:

Blazquez A, Guillén JJ, Colomé C, Coll O, Vassena R, Vernaeve V. Empty follicle syndrome prevalence and management in oocyte donors. Hum Reprod 2014;29:2221–2227.

Blighe K, Rana S, Lewis M. EnhancedVolcano: Publication-ready volcano plots with enhanced colouring and labeling. R package version 2019;1.:

Brower M, Wang E, Hill D, Surrey M, Danzer H, Pisarska MD. The effect of low body mass index (BMI) on ooctye quality in IVF cycles. Fertility and Sterility [Internet] 2013;100:S494.

Burkhardt S, Borsos M, Szydlowska A, Godwin J, Williams SA, Cohen PE, Hirota T, Saitou M, Tachibana-Konwalski K. Chromosome Cohesion Established by Rec8-Cohesin in Fetal Oocytes Is Maintained without Detectable Turnover in Oocytes Arrested for Months in Mice. Curr Biol 2016;26:678–685.

Butler A, Hoffman P, Smibert P, Papalexi E, Satija R. Integrating single-cell transcriptomic data across different conditions, technologies, and species. Nat Biotechnol 2018;36:411–420.

Chamani IJ, Keefe DL. Epigenetics and Female Reproductive Aging. Front Endocrinol 2019;10:473.

Chen H, Boutros PC. VennDiagram: a package for the generation of highly-customizable Venn and Euler diagrams in R. BMC Bioinformatics [Internet] 2011;12.:

Clarke HJ. Growth and Meiotic Maturation of Mammalian Oocytes: An Overview. In Skinner MK, editor. Encyclopedia of Reproduction (Second Edition) 2018;, p. 144–152. Academic Press: Oxford.

De La Fuente R. Chromatin Modifications During Mammalian Oocyte Growth and Meiotic Maturation. In Skinner MK, editor. Encyclopedia of Reproduction (Second Edition) 2018;, p. 183–189. Academic Press: Oxford.

Dobin A, Davis CA, Schlesinger F, Drenkow J, Zaleski C, Jha S, Batut P, Chaisson M, Gingeras TR. STAR: ultrafast universal RNA-seq aligner. Bioinformatics 2013;29:15–21.

Finak G, McDavid A, Yajima M, Deng J, Gersuk V, Shalek AK, Slichter CK, Miller HW, Juliana McElrath M, Prlic M, et al. MAST: a flexible statistical framework for assessing transcriptional changes and characterizing heterogeneity in single-cell RNA sequencing data. Genome Biol [Internet] 2015;16.: BioMed Central.

Fragouli E, Bianchi V, Patrizio P, Obradors A, Huang Z, Borini A, Delhanty JDA, Wells D. Transcriptomic profiling of human oocytes: association of meiotic aneuploidy and altered oocyte gene expression. Mol Hum Reprod 2010;16:570–582.

Ge Z-J, Schatten H, Zhang C-L, Sun Q-Y. Oocyte ageing and epigenetics. Reproduction 2015;149:R103–R114.

Grøndahl ML, Borup R, Vikeså J, Ernst E, Andersen CY, Lykke-Hartmann K. The dormant and the fully competent oocyte: comparing the transcriptome of human oocytes from primordial follicles and in metaphase II. Mol Hum Reprod 2013;19:600–617.

Grøndahl ML, Yding Andersen C, Bogstad J, Nielsen FC, Meinertz H, Borup R. Gene expression profiles of single human mature oocytes in relation to age. Hum Reprod 2010;25:957–968.

Gruhn JR, Zielinska AP, Shukla V, Blanshard R, Capalbo A, Cimadomo D, Nikiforov D, Chan AC-H, Newnham LJ, Vogel I, et al. Chromosome errors in human eggs shape natural fertility over reproductive life span. Science 2019;365:1466–1469.

Guillaumet-Adkins A, Rodríguez-Esteban G, Mereu E, Mendez-Lago M, Jaitin DA, Villanueva A, Vidal A, Martinez-Marti A, Felip E, Vivancos A, et al. Single-cell transcriptome conservation in cryopreserved cells and tissues. Genome Biol 2017;18:45.

Hafemeister C, Satija R. Normalization and variance stabilization of single-cell RNA-seq data using regularized negative binomial regression. Genome Biol 2019;20:296.

Hendrickson PG, Doráis JA, Grow EJ, Whiddon JL, Lim J-W, Wike CL, Weaver BD, Pflueger C, Emery BR, Wilcox AL, et al. Conserved roles of mouse DUX and human DUX4 in activating cleavage-stage genes and MERVL/HERVL retrotransposons. Nat Genet 2017;49:925–934.

Hoffman GE, Le WW, Entezam A, Otsuka N, Tong Z-B, Nelson L, Flaws JA, McDonald JH, Jafar S, Usdin K. Ovarian abnormalities in a mouse model of fragile X primary ovarian insufficiency. J Histochem Cytochem 2012;60:439–456.

Howles CM, Kim C-H, Elder K. Treatment strategies in assisted reproduction for women of advanced maternal age. Int Surg 2006;91:S37–S54.

Huynh-Thu VA, Irrthum A, Wehenkel L, Geurts P. Inferring regulatory networks from expression data using tree-based methods. PLoS One [Internet] 2010;5.:

Hwang G, Sun F, O’Brien M, Eppig JJ, Handel MA, Jordan PW. SMC5/6 is required for the formation of segregation-competent bivalent chromosomes during meiosis I in mouse oocytes. Development 2017;144:1648–1660.

Jalkanen AL, Coleman SJ, Wilusz J. Determinants and implications of mRNA poly(A) tail size--does this protein make my tail look big? Semin Cell Dev Biol 2014;34:24–32.

Labrecque R, Sirard M-A. The study of mammalian oocyte competence by transcriptome analysis: progress and challenges. Molecular Human Reproduction [Internet] 2014;20:103–116.

Lambert SA, Jolma A, Campitelli LF, Das PK, Yin Y, Albu M, Chen X, Taipale J, Hughes TR, Weirauch MT. The Human Transcription Factors. Cell 2018;175:598–599.

Liguori I, Russo G, Curcio F, Bulli G, Aran L, Della-Morte D, Gargiulo G, Testa G, Cacciatore F, Bonaduce D, et al. Oxidative stress, aging, and diseases. Clin Interv Aging 2018;13:757. Dove Press.

Li J-Y, Liu Y-F, Xu H-Y, Zhang J-Y, Lv P-P, Liu M-E, Ying Y-Y, Qian Y-Q, Li K, Li C, et al. Basonuclin 1 deficiency causes testicular premature aging: BNC1 cooperates with TAF7L to regulate spermatogenesis. J Mol Cell Biol 2020;12:71–83.

Lim J, Luderer U. Oxidative damage increases and antioxidant gene expression decreases with aging in the mouse ovary. Biol Reprod 2011;84:775–782.

Lu C, Lin L, Tan H, Wu H, Sherman SL, Gao F, Jin P, Chen D. Fragile X premutation RNA is sufficient to cause primary ovarian insufficiency in mice. Hum Mol Genet 2012;21:5039–5047.

Lu X, Göke J, Sachs F, Jacques P-É, Liang H, Feng B, Bourque G, Bubulya PA, Ng H-H. SON connects the splicing-regulatory network with pluripotency in human embryonic stem cells. Nat Cell Biol 2013;15:1141–1152.

Lu X, Ng H-H, Bubulya PA. The role of SON in splicing, development, and disease. Wiley Interdiscip Rev RNA 2014;5:637–646.

Machtinger R, Combelles CMH, Missmer SA, Correia KF, Fox JH, Racowsky C. The association between severe obesity and characteristics of failed fertilized oocytes. Hum Reprod 2012;27:3198–3207.

Ma J, Zeng F, Schultz RM, Tseng H. Basonuclin: a novel mammalian maternal-effect gene. Development 2006;133:2053–2062.

Matthews TJ, Hamilton BE. Delayed childbearing: more women are having their first child later in life. NCHS Data Brief 2009;1–8.

May-Panloup P, Boucret L, Chao de la Barca J-M, Desquiret-Dumas V, Ferré-L’Hotellier V, Morinière C, Descamps P, Procaccio V, Reynier P. Ovarian ageing: the role of mitochondria in oocytes and follicles. Hum Reprod Update 2016;22:725–743.

Micallef L, Rodgers P. eulerAPE: drawing area-proportional 3-Venn diagrams using ellipses. PLoS One 2014;9:e101717.

Nagaoka SI, Hassold TJ, Hunt PA. Human aneuploidy: mechanisms and new insights into an age-old problem. Nat Rev Genet 2012;13:493–504.

Nakagawa S, FitzHarris G. Intrinsically Defective Microtubule Dynamics Contribute to Age-Related Chromosome Segregation Errors in Mouse Oocyte Meiosis-I. Curr Biol 2017;27:1040–1047.

Namgoong S, Kim N-H. Meiotic spindle formation in mammalian oocytes: implications for human infertility. Biol Reprod 2018;98:153–161.

Olivennes F, Fanchin R, Bouchard P, Taieb J, Frydman R. Triggering of ovulation by a gonadotropin-releasing hormone (GnRH) agonist in patients pretreated with a GnRH antagonist. Fertil Steril 1996;66:151–153.

Pan H, Ma P, Zhu W, Schultz RM. Age-associated increase in aneuploidy and changes in gene expression in mouse eggs. Dev Biol 2008;316:397–407.

Papasaikas P, Tejedor JR, Vigevani L, Valcárcel J. Functional splicing network reveals extensive regulatory potential of the core spliceosomal machinery. Mol Cell 2015;57:7–22.

Picelli S, Björklund åK, Faridani OR, Sagasser S, Winberg G, Sandberg R. Smart-seq2 for sensitive full-length transcriptome profiling in single cells. Nat Methods 2013;10:1096–1098.

Pijuan-Sala B, Guibentif C, Göttgens B. Single-cell transcriptional profiling: a window into embryonic cell-type specification. Nat Rev Mol Cell Biol 2018;19:399–412.

Prochazkova B, Komrskova P, Kubelka M. CPEB2 Is Necessary for Proper Porcine Meiotic Maturation and Embryonic Development. Int J Mol Sci [Internet] 2018;19.:

Reyes JM, Silva E, Chitwood JL, Schoolcraft WB, Krisher RL, Ross PJ. Differing molecular response of young and advanced maternal age human oocytes to IVM. Human Reproduction [Internet] 2017;32:2199–2208.

Saltzman AL, Pan Q, Blencowe BJ. Regulation of alternative splicing by the core spliceosomal machinery. Genes Dev 2011;25:373–384.

Saskova A, Solc P, Baran V, Kubelka M, Schultz RM, Motlik J. Aurora kinase A controls meiosis I progression in mouse oocytes. Cell Cycle 2008;7:2368–2376.

Shah DK, Missmer SA, Berry KF, Racowsky C, Ginsburg ES. Effect of obesity on oocyte and embryo quality in women undergoing in vitro fertilization. Obstet Gynecol 2011;118:63–70.

Shannon P, Markiel A, Ozier O, Baliga NS, Wang JT, Ramage D, Amin N, Schwikowski B, Ideker T. Cytoscape: a software environment for integrated models of biomolecular interaction networks. Genome Res 2003;13:2498–2504.

Sha Q-Q, Yu J-L, Guo J-X, Dai X-X, Jiang J-C, Zhang Y-L, Yu C, Ji S-Y, Jiang Y, Zhang S-Y, et al. CNOT6L couples the selective degradation of maternal transcripts to meiotic cell cycle progression in mouse oocyte. EMBO J [Internet] 2018;37.:

Sherman SL, Curnow EC, Easley CA, Jin P, Hukema RK, Tejada MI, Willemsen R, Usdin K. Use of model systems to understand the etiology of fragile X-associated primary ovarian insufficiency (FXPOI). J Neurodev Disord 2014;6:26.

Shirai Y-T, Suzuki T, Morita M, Takahashi A, Yamamoto T. Multifunctional roles of the mammalian CCR4-NOT complex in physiological phenomena. Front Genet 2014;5:286.

Smoak EM, Stein P, Schultz RM, Lampson MA, Black BE. Long-Term Retention of CENP-A Nucleosomes in Mammalian Oocytes Underpins Transgenerational Inheritance of Centromere Identity. Curr Biol 2016;26:1110–1116.

Solc P, Baran V, Mayer A, Bohmova T, Panenkova-Havlova G, Saskova A, Schultz RM, Motlik J. Aurora Kinase A Drives MTOC Biogenesis but Does Not Trigger Resumption of Meiosis in Mouse Oocytes Matured In Vivo1. Biology of Reproduction [Internet] 2012;87.:

Steuerwald NM, Bermúdez MG, Wells D, Munné S, Cohen J. Maternal age-related differential global expression profiles observed in human oocytes. Reproductive BioMedicine Online [Internet] 2007;14:700–708.

Stuart T, Butler A, Hoffman P, Hafemeister C, Papalexi E, Mauck WM 3rd, Hao Y, Stoeckius M, Smibert P, Satija R. Comprehensive Integration of Single-Cell Data. Cell 2019;177:1888–1902.e21.

Sun S, Liu J, Zhao M, Han Y, Chen P, Mo Q, Wang B, Chen G, Fang Y, Tian Y, et al. Loss of the novel mitochondrial protein FAM210B promotes metastasis via PDK4-dependent metabolic reprogramming. Cell Death Dis 2017;8:e2870.–e2870. Nature Publishing Group.

Susor A, Kubelka M. Translational Regulation in the Mammalian Oocyte. Results and Problems in Cell Differentiation [Internet] 2017;257–295Available from: http://dx.doi.org/10.1007/978-3-319-60855-6_12.

Su Y-Q, Sugiura K, Woo Y, Wigglesworth K, Kamdar S, Affourtit J, Eppig JJ. Selective degradation of transcripts during meiotic maturation of mouse oocytes. Dev Biol 2007;302:104–117.

Svensson V, Vento-Tormo R, Teichmann SA. Exponential scaling of single-cell RNA-seq in the past decade. Nat Protoc 2018;13:599–604.

Swartz SZ, McKay LS, Su K-C, Bury L, Padeganeh A, Maddox PS, Knouse KA, Cheeseman IM. Quiescent Cells Actively Replenish CENP-A Nucleosomes to Maintain Centromere Identity and Proliferative Potential. Dev Cell 2019;51:35–48.e7.

Tachibana-Konwalski K, Godwin J, Weyden L van der, Champion L, Kudo NR, Adams DJ, Nasmyth K. Rec8-containing cohesin maintains bivalents without turnover during the growing phase of mouse oocytes. Genes Dev 2010;24:2505–2516.

Tacutu R, Thornton D, Johnson E, Budovsky A, Barardo D, Craig T, Diana E, Lehmann G, Toren D, Wang J, et al. Human Ageing Genomic Resources: new and updated databases. Nucleic Acids Res 2018;46:D1083–D1090.

Toth A, Jessberger R. Oogenesis: Ageing Oocyte Chromosomes Rely on Amazing Protein Stability. Curr Biol 2016;26:R329–R331.

Vieux K-F, Clarke HJ. CNOT6 regulates a novel pattern of mRNA deadenylation during oocyte meiotic maturation. Sci Rep 2018;8:6812.

Virant-Klun I, Leicht S, Hughes C, Krijgsveld J. Identification of Maturation-Specific Proteins by Single-Cell Proteomics of Human Oocytes. Mol Cell Proteomics 2016;15:2616–2627.

Wang S, Zheng Y, Li J, Yu Y, Zhang W, Song M, Liu Z, Min Z, Hu H, Jing Y, et al. Single-Cell Transcriptomic Atlas of Primate Ovarian Aging. Cell 2020;180:585–600.e19.

Wickham H. ggplot2: Elegant Graphics for Data Analysis. 2016; Springer.

Wilkinson L. Exact and approximate area-proportional circular Venn and Euler diagrams. IEEE Trans Vis Comput Graph 2012;18:321–331.

Yan C, Wang P, DeMayo J, DeMayo FJ, Elvin JA, Carino C, Prasad SV, Skinner SS, Dunbar BS, Dube JL, et al. Synergistic roles of bone morphogenetic protein 15 and growth differentiation factor 9 in ovarian function. Mol Endocrinol 2001;15:854–866.

Yu G, Wang L-G, Han Y, He Q-Y. clusterProfiler: an R Package for Comparing Biological Themes Among Gene Clusters. OMICS: A Journal of Integrative Biology [Internet] 2012;16:284–287.

Zhang D, Liu Y, Zhang Z, Lv P, Liu Y, Li J, Wu Y, Zhang R, Huang Y, Xu G, et al. Basonuclin 1 deficiency is a cause of primary ovarian insufficiency. Hum Mol Genet 2018;27:3787–3800.

Zhang J-J, Liu X, Chen L, Zhang S, Zhang X, Hao C, Miao Y-L. Advanced maternal age alters expression of maternal effect genes that are essential for human oocyte quality. Aging 2020;12:3950–3961.

Zhang S, Wang J, Tseng H. Basonuclin regulates a subset of ribosomal RNA genes in HaCaT cells. PLoS One 2007;2:e902.

Zielinska AP, Bellou E, Sharma N, Frombach A-S, Seres KB, Gruhn JR, Blayney M, Eckel H, Moltrecht R, Elder K, et al. Meiotic Kinetochores Fragment into Multiple Lobes upon Cohesin Loss in Aging Eggs. Curr Biol 2019;29:3749–3765.e7.

