## Supplementary Fig.1-5 + Table Legends for "Single cell analysis reveals the impact of age and maturation stage on the human oocyte transcriptome"

### Supplementary data

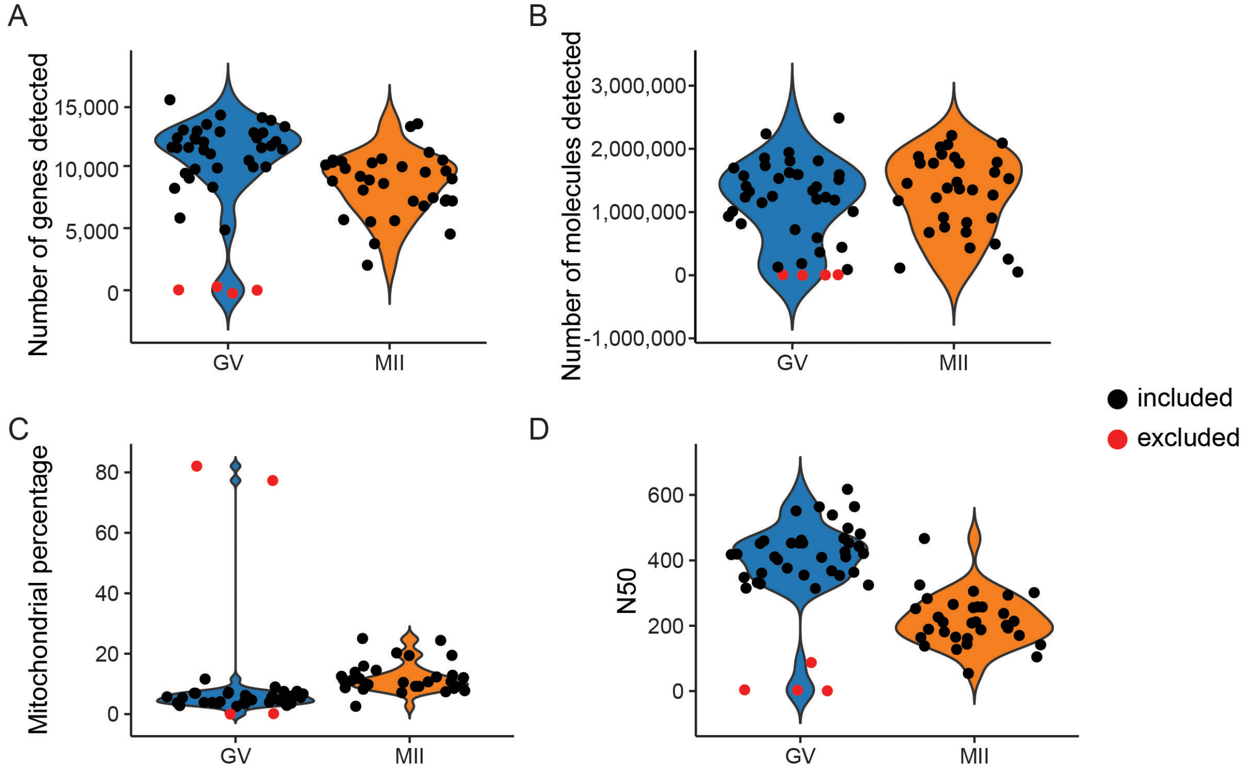


**Supplementary Figure S1. Quality control applied to the dataset.** Oocytes presenting any of the following characteristics (shown in red) were considered of low quality and were removed from the analysis: over 30% of mitochondrial percentage, less than 1000 genes and under 10000 reads.

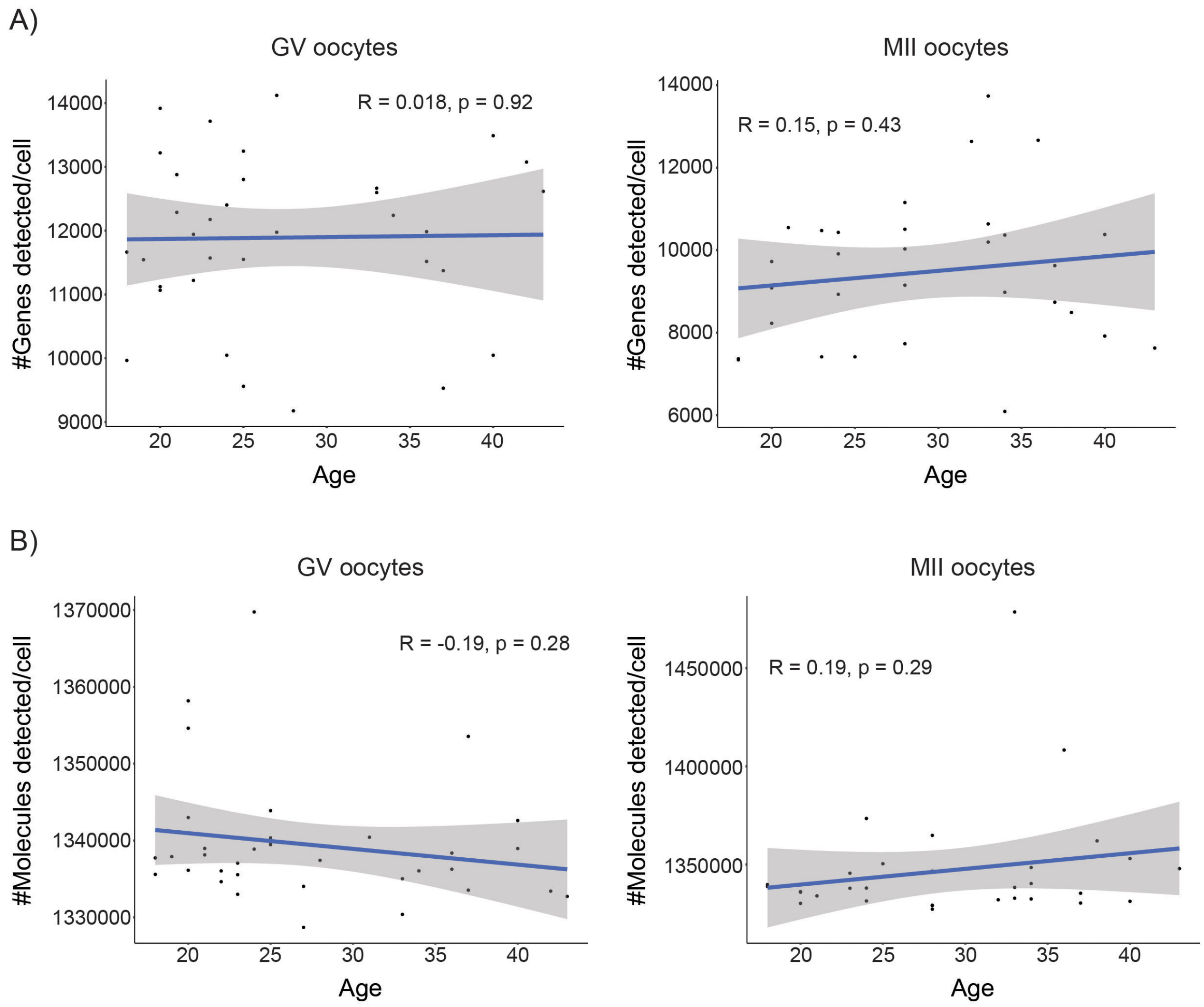


**Supplementary Figure S2. Brief exploration of the sample set.** Neither the total number of genes **(A)** nor the total number of RNA molecules **(B)** detected in both germinal vesicle stage oocytes and *in vitro* matured metaphase II stage oocytes varied significantly with age.

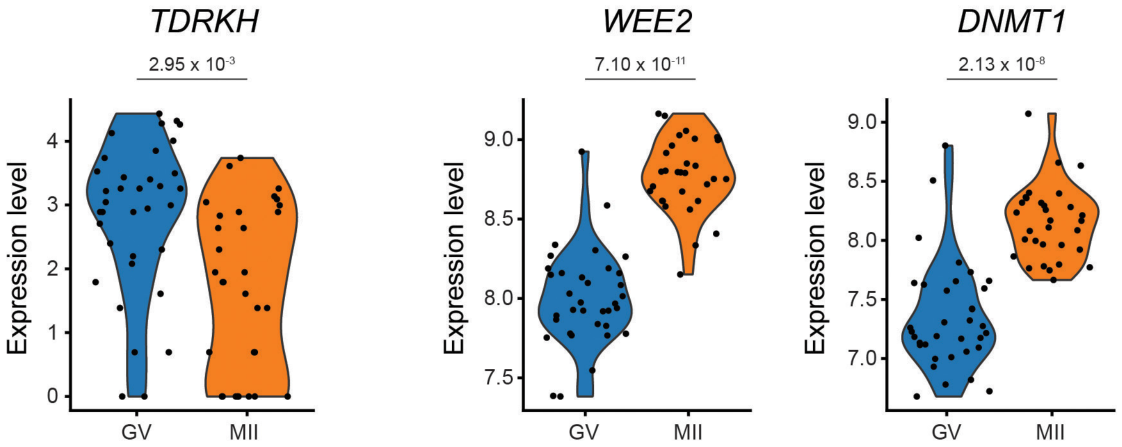


**Supplementary Figure S3. RNA abundance of published protein markers** (Virant-Klun *et al.*, 2016) of GV (TDRKH) and MII stage (WEE2, DNMT1) oocytes in our dataset.

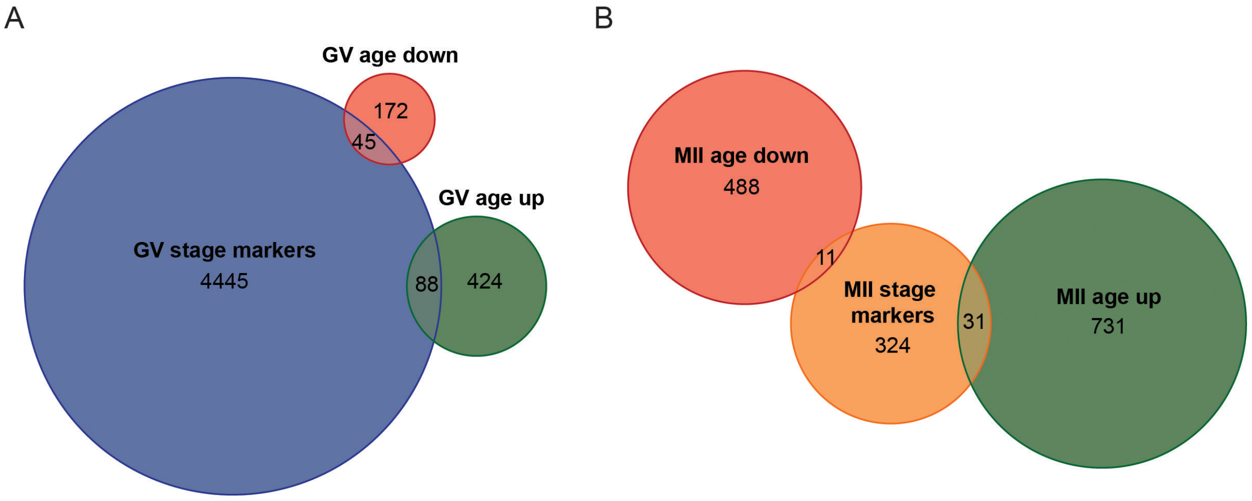


**Supplementary Figure S4.** Intersection of maturation stage markers with genes changing in transcript abundance with age. **(A)** Venn diagram of overlap of GV stage markers with genes, for which RNA levels increase (GV age up) or decrease (GV age down) with age. (**B)** Overlap between IVM-MII stage markers and genes changing in transcript abundance with age in IVM-MII oocytes.

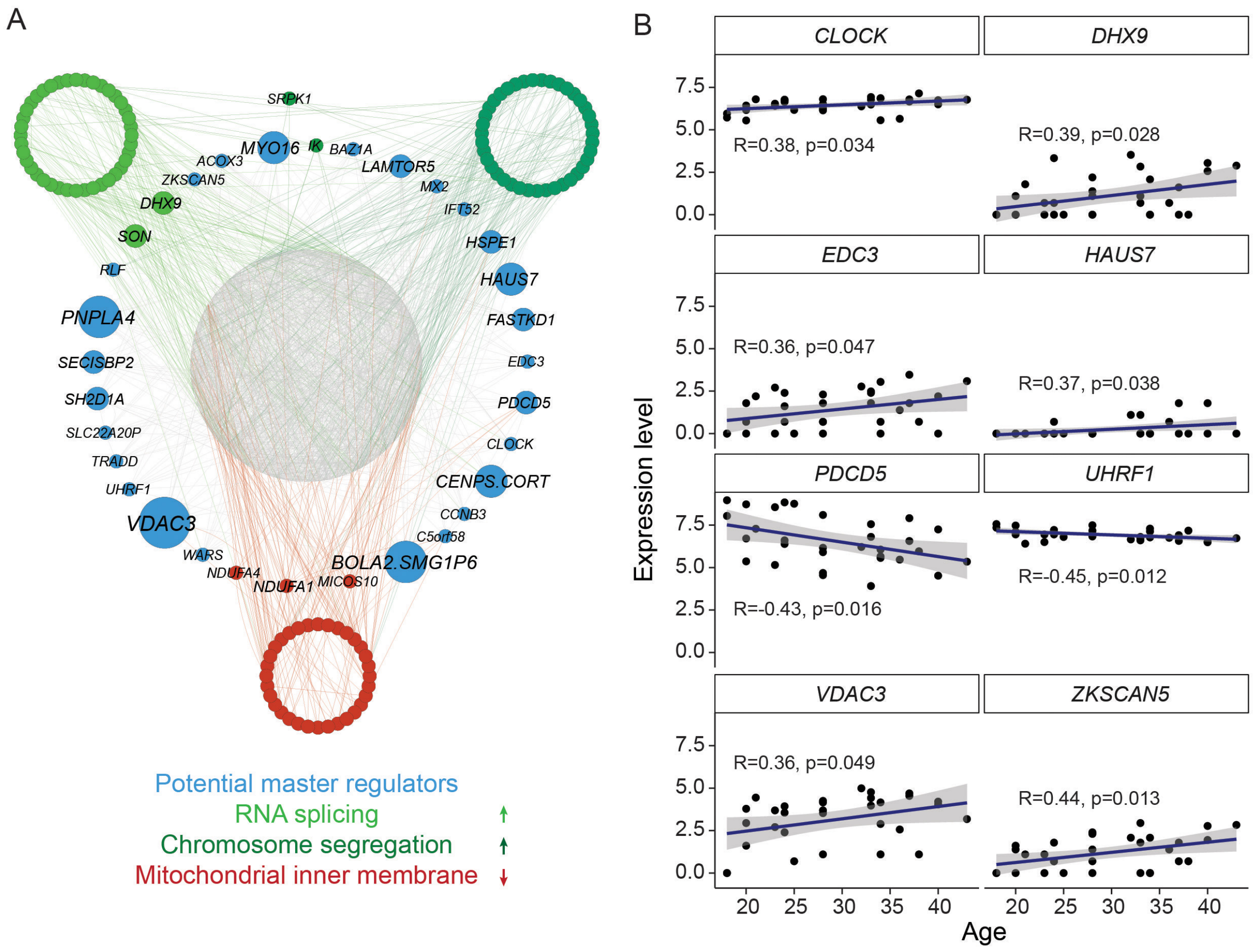


**Supplementary Figure S5. Gene regulatory network analysis. (A)** Cytoscape plots from the top 2,500 regulatory links among genes found to correlate with age in IVM-MII oocytes. In that case, all the DEG genes were given as an input to Genie3 to be used as potential regulators of the network. In green are represented the genes belonging to two of the GO terms enriched in genes whose RNA levels correlate positively with age, "RNA splicing" (light green) and "chromosome segregation" (dark green). *SRPK1* and *IK* belong to both of these GO terms, and therefore they are plotted in between. In red are represented genes belonging to "mitochondrial inner membrane", which appears as an enriched GO term in the list of genes whose RNA levels correlate negatively with age. **(B)** Plots showing how the expression levels of the potential master regulators of the network vary with age.

### Supplementary table legends

**Supplementary Table 1**: Individual characteristics of women participating in the study and their sample contribution

**Supplementary Table 2.** Maturation stage (GV and IVM-MII) markers. For each marker, the average expression level, standard deviation, p-value and average fold change (in relation to the other stage) are shown.

**Supplementary Table 3.** Gene Ontology results for the markers of each maturation stage. The ontology category, GO ID and description, as well as the genes present in each gene ontology term are included.

**Supplementary Table 4.** Genes changing in RNA levels with age. The first tab contains a list with the genes which correlate with age. The p-values and Pearson correlation values are shown. The following tabs contain the filtered genes: genes that increase expression levels with age in GVs, genes that decrease expression levels with age in GVs, genes that increase expression levels with age in IVM-MIIs and genes that decrease expression levels with age in IVM-MIIs.

**Supplementary Table 5.** Gene Ontology results for the genes that correlate with age in IVM-MII stage oocytes. First tab: RNAs that increase with age. Second Tab: RNAs that decrease with age.

**Supplementary Table 6.** Link lists obtained from the Genie3 network analysis of age related expression changes. The first column contains the regulatory genes, the second column contains the target gene and the third column contains the weight, a value without statistical significance meaning and that serves to rank the regulatory links. First tab: Transcription factors (TF) selected as potential regulators. Second Tab: Any gene as potential regulator (no-TF).

**Supplementary Table 7.** Genes changing in RNA levels with BMI. The first tab contains a list with the genes which correlate with BMI. The p-values and Pearson correlation values are shown. The following tabs contain the filtered genes: genes that increase expression levels with BMI in GVs, genes that decrease expression levels with BMI in GVs, genes that increase expression levels with BMI in IVM-MIIs and genes that decrease expression levels with BMI in IVM-MIIs.

**Supplementary Table 8.** Gene ontology results for genes that correlate with BMI in GV stage oocytes. First tab: RNAs that increase with BMI. Second Tab: RNAs that decrease with BMI.
