## Supplementary Table 1 for "Single cell analysis reveals the impact of age and maturation stage on the human oocyte transcriptome"

**Supplementary Table 1**: Individual characteristics of women participating in the study and their sample contribution; women ordered by age at participation. AFC: Antral Follicular Count; BMI: Body Mass Index; GV: Germinal Vesicle stage oocytes included; IVM-MII: MII after in vitro maturation oocytes included; P: patient; D: donor.

| WOMAN | AGE (years) | AFC (n) | BMI (kg/m2) | Patient or Donor | GV (n) | IVM-MII (n) |
|---|---|---|---|---|---|---|
| 1 | 18 | 46 | 28 | D | 2 | 2 |
| 2 | 19 | 15 | 20 | D | 1 | 0 |
| 7 | 20 | 27 | 17 | D | 1 | 0 |
| 3 | 20 | 29 | 19 | D | 1 | 1 |
| 4 | 20 | 22 | 20 | D | 2 | 2 |
| 6 | 20 | 15 | 23 | D | 1 | 0 |
| 5 | 20 | 10 | 25 | D | 1 | 0 |
| 9 | 21 | 10 | 19 | D | 1 | 1 |
| 8 | 21 | 28 | 21 | D | 1 | 0 |
| 10 | 22 | 22 | 17 | D | 2 | 0 |
| 11 | 23 | 24 | 27 | D | 1 | 1 |
| 12 | 23 | 35 | 27 | D | 2 | 1 |
| 13 | 24 | 31 | 20 | D | 1 | 1 |
| 14 | 24 | 46 | 22 | D | 2 | 1 |
| 15 | 24 | 23 | 26 | D | 0 | 1 |
| 17 | 25 | 25 | 24 | D | 2 | 0 |
| 16 | 25 | 22 | 27 | D | 2 | 1 |
| 18 | 27 | 30 | 23 | D | 2 | 0 |
| 19 | 28 | 35 | 21 | D | 1 | 1 |
| 21 | 28 | 19 | 21 | D | 0 | 2 |
| 20 | 28 | 28 | 23 | D | 0 | 2 |
| 22 | 31 | 5 | 24 | P | 1 | 0 |
| 23 | 32 | 34 | 25 | P | 0 | 1 |
| 24 | 33 | 11 | 23 | D | 0 | 1 |
| 26 | 33 | 17 | 26 | D | 1 | 1 |
| 25 | 33 | 25 | 29 | D | 1 | 1 |
| 28 | 34 | 17 | 26 | D | 1 | 1 |
| 27 | 34 | 6 | 28 | P | 0 | 2 |
| 30 | 36 | 10 | 19 | P | 1 | 0 |
| 29 | 36 | >25 | 30 | P | 2 | 1 |
| 31 | 37 | 32 | 23 | P | 2 | 2 |
| 32 | 38 | 25 | 23 | P | 0 | 1 |
| 33 | 40 | 4 | 20 | P | 1 | 2 |
| 34 | 40 | 17 | 32 | P | 1 | 1 |
| 35 | 42 | 29 | 26 | P | 1 | 0 |
| 36 | 43 | 7 | 24 | P | 1 | 0 |
| 37 | 43 | 16 | 25 | P | 1 | 1 |
| TOTAL | | | | | 40 | 32 |
